# Human intestinal epithelial cells can internalize luminal fungi via LC3-associated phagocytosis

**DOI:** 10.1101/2022.12.14.520372

**Authors:** Sarit Cohen-Kedar, Efrat Shaham Barda, Keren Masha Rabinowitz, Danielle Keizer, Hannan Abu-Taha, Shoshana Schwartz, Kawsar Kaboub, Liran Baram, Eran Sadot, Ian White, Nir Wasserberg, Meirav Wolff-Bar, Adva Levy-Barda, Iris Dotan

## Abstract

Intestinal epithelial cells (IECs) are the first to encounter luminal microorganisms and actively participate in intestinal immunity. We reported that IECs express the β-glucan receptor Dectin-1, and respond to commensal fungi and β-glucans. In phagocytes, Dectin-1 mediates LC3 associated phagocytosis (LAP) utilizing autophagy components to process extracellular cargo. Dectin-1 can mediate phagocytosis of β-glucan-containing particles by non-phagocytic cells. We aimed to determine whether human IECs phagocytose β-glucan-containing fungal particles via LAP. Zymosan (β-glucan particle) and Heat-killed and UV inactivated *C. albicans* were phagocytosed by monolayers of human colonic (n=18) and ileal (n=4) organoids and IEC lines. LAP was identified by LC3 and Rubicon recruitment to phagosomes and lysosomal processing of internalized particles was demonstrated by co-localization with lysosomal dyes and LAMP2. Phagocytosis was significantly diminished by blockade of Dectin-1, actin polymerization and NAPDH oxidases. Our results show that human IECs sense luminal fungal particles and internalize them via LAP. This novel mechanism of luminal sampling suggests that IECs may contribute to the maintenance of mucosal tolerance towards commensal fungi.

## Introduction

Intestinal epithelial cells (IECs) stand in the frontline of the largest mucosal surface in the human body. As such, IECs are in constant interaction with luminal microorganisms and dietary molecules, as well as with immune cells in the lamina propria beneath them. Besides being a physical barrier, IECs function as innate immune cells, sensing and actively responding to luminal microbiota (Allaire et al., 2018; Gunther and Seyfert, 2018; Soderholm and Pedicord, 2019). However, the role of human IECs in host tolerance towards commensal fungi and their contribution to shaping host immune response remain obscure.

We previously reported that human IECs express the β-glucan receptor Dectin-1 (Cohen-Kedar et al., 2014), a central C-type-lectin-receptor (CLR) involved in fungal recognition and immune response (Del Fresno et al., 2018; Höft et al., 2020; Mata-Martínez et al., 2022; Nikolakopoulou et al., 2020; Salazar and Brown, 2018). We further demonstrated that IECs were directly activated by cell wall components of commensal fungi via Dectin-1. However, pro-inflammatory cytokine secretion occurring in response to β-glucan was silenced when whole fungi were sensed (Cohen-Kedar et al., 2021) suggesting epithelial tolerance to commensals.

We therefore sought a physiological homeostatic outcome of fungal recognition by IECs. Dectin-1 functions in phagocytosis of non-opsonized fungi by professional phagocytes (Brown and Gordon, 2001; Mata-Martínez et al., 2022; Taylor et al., 2007). As Dectin-1 expression allows β-glucan-dependent phagocytosis by non-phagocytes (Brown and Gordon, 2001; Gantner et al., 2005; Herre et al., 2004), we asked whether IECs can phagocytose β -glucan containing fungal particles, and by which mechanism.

Here we provide evidence supporting a novel mechanism of interaction between IECs and commensal fungi at the intestinal mucosa, where β-glucan and *C. albicans* were phagocytosed by human IECs in a Dectin-1 dependent and spleen tyrosine kinase (Syk) independent manner leading to LC3 associated phagocytosis (LAP) and lysosomal degradation.

## Results

### Zymosan uptake by IECs is dependent on actin-polymerization

To assess phagocytosis in IECs we chose zymosan, a particulate β-glucan rich, cell wall extract of *Saccharomyces cerevisiae* which is commensal in the human gut. Hence, zymosan is highly applicable as a representative of luminal fungal species in the vicinity of mucosal surfaces. In addition, zymosan is widely used to study phagocytosis by professional phagocytes and the pHrodo-red label of zymosan, that turns fluorescent intracellularly, is indicative of internalized particles in living cells (Lindner et al., 2020). Using the human epithelial cell lines SW480, HCT116 and Caco-2 we detected cellular uptake of pHrodo-red zymosan where some cells presented multiple internalized particles as well as fragmented zymosan indicative of intracellular processing (Fig. 1A and Supp. Fig. S1A, S2A respectively), suggesting that phagocytosis of β-glucan expressing particles is common in human IECs. We detected cellular internalization of pHrodo-red zymosan in a few cells as early as 3 hours of incubation and phagocytosis was clearly observed following over-night incubation where up to 20% of the cells were zymosan positive (Fig. 1B, D).

**Figure 1:**
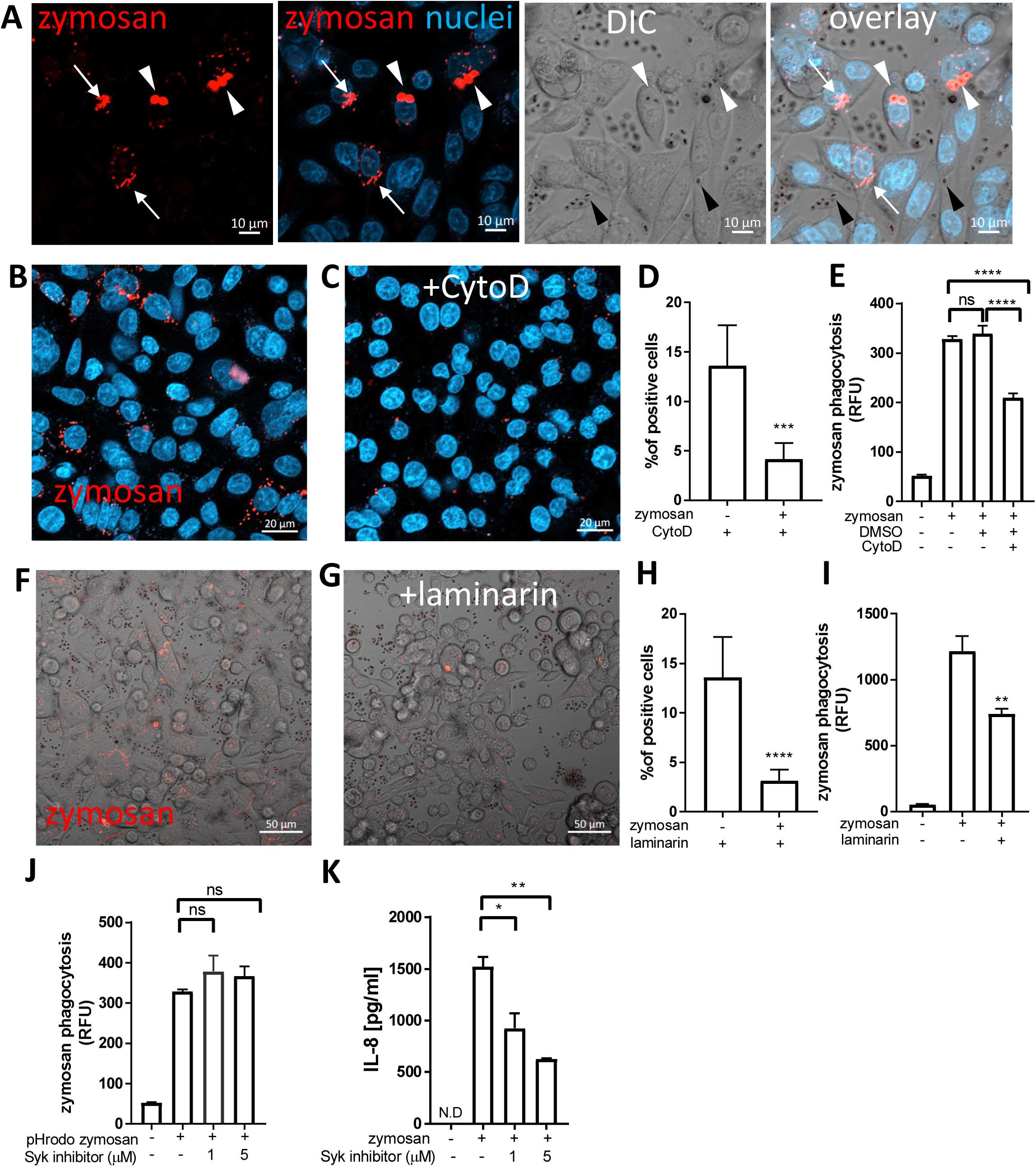
Uptake of zymosan by human intestinal epithelial cells. (A) SW480 cells were seeded on glass-bottom chambers as indicated in Methods, and fed over-night with pHrodo-red zymosan (zymosan, red) and counter stained with Hoechst 33342 (blue) prior to confocal live imaging. White arrowheads – intracellular red fluorescent zymosan, Black arrowheads - extracellular intact zymosan, arrow-intracellular fragmented zymosan. Original magnification x20, scale bar 10 µm. (B-E) Zymosan uptake is sensitive to cytochalasin-D. (B, C) SW480 were treated as in A, in the absence (B) or presence (C) of cytochalasin-D (CytoD, 10 µM). Scale bar 20 µm. Wider fields of the images are shown is Supp. Fig. S3. (D) Phagocytosis was quantified using imageJ as the percentage of red-fluorescence positive cells in at least 4 randomly taken fields as described in Methods. Data is representative of three independent experiments performed. ***p≤0.001, Unpaired t-test vs. no inhibitor. (E) SW480 cells were seeded in 96 well plate, treated as in (B-C) as well as with the vehicle (DMSO, 1:1000) in triplicate wells, and phagocytosis was assessed as the relative fluorescence (RFU) by a microplate reader. Data are shown as mean ± SD of biological triplicates from a representative of three independent experiments performed. ns-non significant ****p<0.0001, One-way ANOVA followed by Tukey multiple comparison test. (F-I) Zymosan uptake depends on Dectin-1. (F,G) SW480 were treated as in A, in the absence (F) or presence (G) of laminarin (1 mg/ml) that was added to the medium 1 hour prior to zymosan. Scale bar 50 µm. (H) Phagocytosis was quantified as in (D). (I) cells were seeded on 96 wells, treated as in (F,G) in triplicate wells, and phagocytosis was analyzed as in (E). (J) Zymosan phagocytosis is resistant to Syk inhibition. SW480 cells were seeded on 96 well plate, in the presence or absence of the Syk inhibitor 574711 (1 and 5 µM), which was added 1 hour prior to the addition of pHrodo-red zymosan. Phagocytosis was assessed as in (E). Data are shown as mean ± SD of biological triplicates from a representative of three independent experiments performed. (K) Zymosan-induced IL-8 secretion is sensitive to Syk inhibitor. Cells seeded on the same 96 well plate were pre-treated with Syk inhibitor as in (J) and stimulated over-night with 100 ug/ml of non-labelled zymosan. Supernatants were assessed for IL-8 by ELISA. Data are shown as mean ± SD of biological duplicates from a representative of three independent experiments performed. N.D-not detected; ns-non significant *p<0.05; **p<0.01; ***p<0.001; ****p<0.0001, One-way ANOVA followed by Tukey multiple comparison test.

To test whether cytoskeleton-mediated engulfment participates in the uptake of zymosan, we applied the actin-polymerization-inhibitor cytochalasin-D. This resulted in attenuated phagocytosis as reflected by a 70% decrease in the number of pHrodo-red zymosan positive SW480 cells (***p=≤0.001, Fig. 1B-D and Supp. Fig. S3), and in the total fluorescence of intracellular pHrodo-red zymosan (**p≤0.01, Fig. 1E).

Similar sensitivity to cytochalasin-D was observed in HCT116 cells (Supp. Fig. S1B-C). As actin mediated engulfment of extracellular particles is a hallmark of phagocytosis these results indicate a genuine zymosan phagocytosis in IECs.

### Zymosan phagocytosis by IECs involves Dectin-1

We have previously demonstrated functional Dectin-1 in IECs (Cohen-Kedar et al., 2014; Cohen-Kedar et al., 2021). Since Dectin-1 mediates phagocytosis in professional phagocytes, we next asked whether it also functions in phagocytosis in IECs. To this end, we used laminarin, a soluble Dectin-1 antagonist, that blocks zymosan and fungal phagocytosis in professional phagocytes (Fuentes et al., 2011; Kyrmizi et al., 2013; Luther et al., 2007; Maneu et al., 2011). Notably, laminarin inhibited zymosan phagocytosis by SW480 and HCT116 cells as indicated by significant decrease in the number of pHrodo-red zymosan positive cells (by 77% ****p≤0.0001, Fig. 1F-H and Supp. Fig. S1B) and the total fluorescence of the intracellular pHrodo-red zymosan (Fig. 1I **p=≤0.01 and S1D ***p=≤0.001). Altogether Dectin-1 dependent zymosan phagocytosis by IEC lines was demonstrated.

### Phagocytosis by IECs is Syk-independent

Syk is a major signaling mediator down-stream Dectin-1 and is involved in Dectin-1 triggered phagocytosis by professional phagocytes (Kyrmizi et al., 2013; Ma et al., 2012). Yet, there are examples where Syk was dispensable for phagocytosis (Herre et al., 2004; Underhill et al., 2005). Syk is activated by commensal fungi and β-glucan and is required for β-glucan-induced cytokine secretion in human IECs (Cohen-Kedar et al., 2014; Cohen-Kedar et al., 2021; Wang et al., 2021). Therefore, we asked whether Syk is essential for zymosan phagocytosis. We found that the Syk inhibitor 574711 [3-(1-methyl-1H-indol-3-yl-methylene)-2-oxo-2,3-dihydro-1H-indole-5-sulfonamide] did not interfere with phagocytosis (Fig. 1J) while its inhibitory activity was indicated by a significant decrease of zymosan-induced IL-8 secretion in the same experiment (60% inhibition by 5 µM of Syk inhibitor, **P≤0.01, Fig. 1K). Further evidence for Syk-independence was obtained with Caco2 cells, which do not express Syk (Cohen-Kedar et al., 2021), but readily phagocytose pHrodo-red zymosan (Supp. Fig. S2). We conclude that in IECs, phagocytosis of zymosan may occur independently of Syk activation.

### Human intestinal organoids can phagocytose zymosan

We next asked whether primary human IECs can phagocytose zymosan. To test this, we used human intestinal organoids generated from ileal and colonic crypts obtained from surgical samples, that were grown as two-dimensional monolayers to facilitate epithelial exposure to large particles (Dutta et al., 2017)(see methods). We assessed phagocytosis in ileal and colonic organoids cultured in expansion medium and then grown for 2-3 additional days in a generic differentiation medium (see methods) prior to pHrodo-red zymosan exposure (Fig. 2A, 2B, Supp. Figs. S4 and S5A). We found pronounced phagocytic activity, which was distinctly higher in organoids grown in differentiation medium compared to those cultured in expansion medium only (Supp. Fig. S5B). Hence, we performed our phagocytosis experiments in differentiation medium throughout this work. Phagocytosis of zymosan was observed by all ileal (n=4) and colonic (n=18, derived from ascending [n=13], transverse [n=2] or from sigmoid colon [n=3]) organoids tested, suggesting that phagocytic capacity of the epithelium is found along the human gastrointestinal tract. Fragmentation of internalized (fluorescent) zymosan suggests its intracellular processing in organoid cells (Fig. 2C and supp. movie. 1 and 2), similarly to our observation in cell lines.

**Figure 2:**
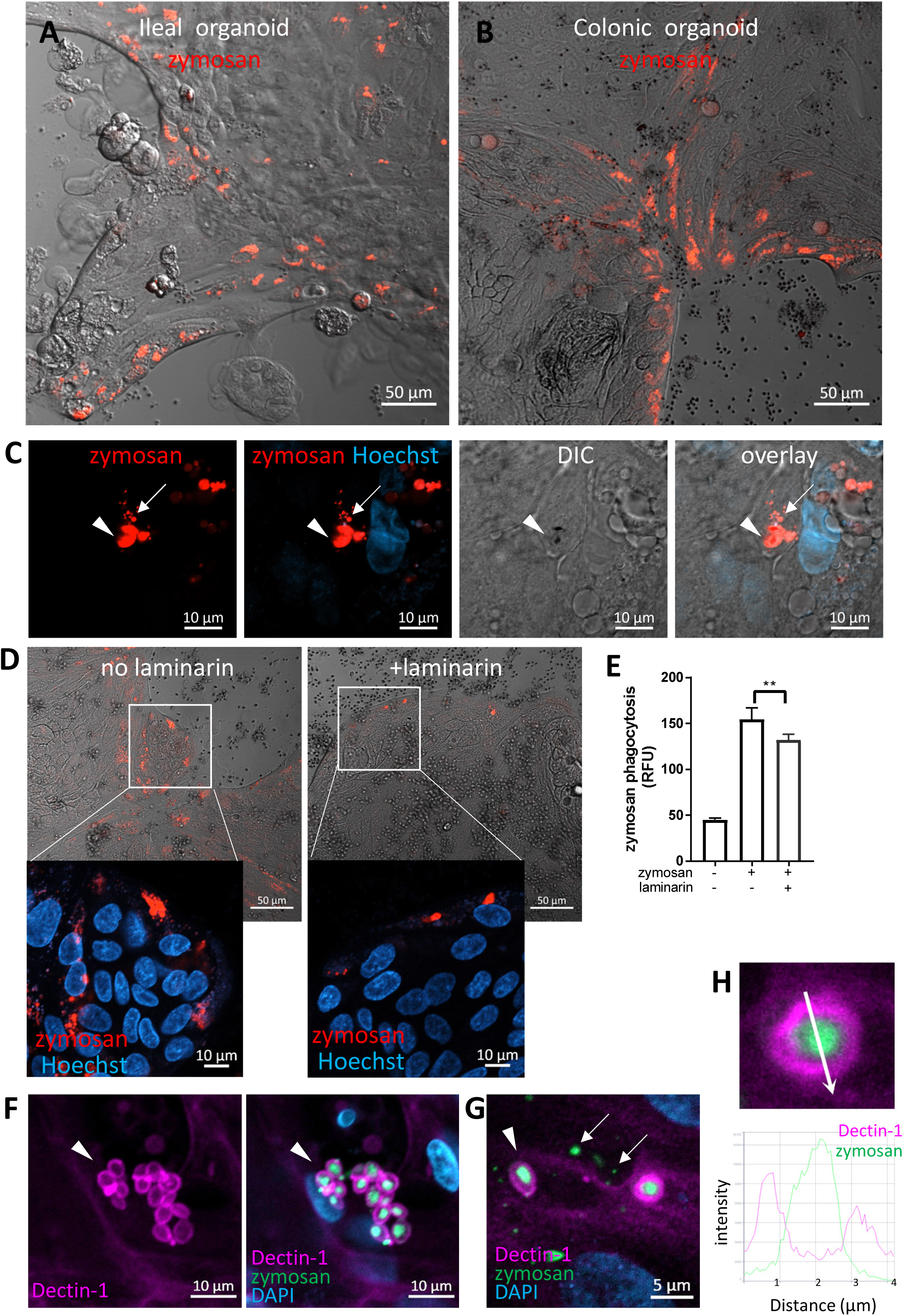
Intestinal organoids uptake zymosan. (A,B) Ileal (A) and colonic (B) organoids were grown as monolayers in expansion medium and let do differentiate for 2 days. pHrodo-red zymosan (red) was added to the medium for 24h. Original magnification x10, scale bar 50 µm. Shown are representative frames from wider fields, presented in Supp. Figs S4 and S5A, from 3-5 randomly acquired scans of two independent experiments of two organoids. (C) Colonic organoids (from a different individual) were treated as in (B), nuclei were stained with Hoechst 33342 (blue) prior to confocal live imaging. Shown a representative frame from a z-stuck analysis. Entire z-stuck movie is shown in Supp. Movie 1. Arrowheads – intracellular fluorescent zymosan, arrows- intracellular fragmented zymosan. Original magnification x63, scale bar 10 µm. (D) Laminarin inhibits zymosan uptake by intestinal organoids. Colonic organoids were grown as in B, in the presence or absence of laminarin (1 mg/ml) that was added to the medium 1 hour prior to zymosan in triplicate wells. Shown are intracellular fluorescent zymosan (red) and the organoid cells (DIC), or zoom-in insets where nuclei were stained with Hoechst 33342 (blue) prior to confocal live imaging. Shown are representative frames from 3-6 random fields imaged from each of triplicate wells. The experiment was repeated with 4 different organoids. Original magnification x20, scale bar 50 and 10 µm. (E) Colonic organoids were seeded in 96 well plate, treated as in (D) in triplicate wells for 48 hours. Phagocytosis was assessed as the relative fluorescence by a microplate reader. Data are shown as mean ± SD of biological triplicates from a representative of two independent experiments performed. ***p<0.001 vs. no inhibitor, Student’s t-test. (F,G) Dectin-1 is recruited to internalized zymosan. Ileal organoids were fed with AF488-zymosan (green) over-night and stained with Dectin-1 antibody (magenta) and DAPI. Original magnification x20 scale bar 10 µm (F) and x63 scale bar 5 µm (G). Arrowheads – intact zymosan, arrows-fragmented zymosan. (H) Fluorescence intensity profile along the arrow of an inset from (G) is shown on the graph.

Notably, different epithelial cell types in the organoid monolayers were (Supp. Fig. S6 and S7) and phagocytosis was observed in goblet as (MUC2^+^) as well as non-goblet (MUC2^-^) IECs (Supp. Fig. S8). This appeared to be more prominent at the periphery of the organoid cultures, which represent the luminal regions of the villus or crypt (Thorne et al., 2018). Interestingly, Dectin-1 was expressed at the peripheral regions of the organoids (Supp. Fig. S9) and at the apical side of the lumen-facing IECs in colonic crypts of frozen sections (Supp. Fig. S10). To test the function of Dectin-1 in phagocytosis, we assessed intracellular pHrodo-red zymosan in organoids in the presence or absence of laminarin. A decrease in intracellular pHrodo-red zymosan in the presence of laminarin (Fig. 2D) was observed by live confocal microscopy. This observation was quantitatively confirmed by a significant decrease (***p≤0.001) in the fluorescence of pHrodo-red zymosan measured by a microplate reader (Fig. 2E). Next, we show that Dectin-1 engulfs intracellular zymosan particles, as verified by fluorescence profile analysis (Fig. 2F-H and Supp. Fig. S11) while it was not detected at fragmented zymosan (Fig. 2G). This implies a specific role for Dectin-1 at the early stages of zymosan recognition and uptake, rather than during intracellular processing. Collectively, our results suggest that human colonic and ileal IECs are able to phagocytose zymosan in a Dectin-1-dependent manner.

### Phagocytosis of *Candida albicans* by IECs

We next asked whether IECs phagocytose fungal particles. *C. albicans*, in its yeast-form, is a frequent commensal in the human gastrointestinal tract (Li et al., 2022; Sokol et al., 2017). The cell-wall inner β-glucan layer is exposed in heat-killed (HK) *C. albicans,* rendering it highly accessible to Dectin-1. Yet, unlike zymosan, HK-*C. albicans* did not induce cytokine secretion by IECs, although it did elicit Syk and ERK activation (Cohen-Kedar et al., 2021). We therefore labeled a commercially available preparation of HK-*C. albicans* strain (ATCC 10231) with Rhodamine-green-X to assess microscopically its uptake by IECs. Colonic and ileal organoids, as well as IEC lines internalized HK-*C. albicans* particles where cellular fragmentation verified their intracellular localization and processing (Fig. 3A, Supp. fig S12A). As observed for zymosan, here too, goblet and non-goblet cells (MUC2^+^ and MUC2^-^ respectively) phagocytosed HK-*C. albicans* (Supp. Fig.S13, Fig.S14). Simultaneous exposure of organoid cultures to HK-*C. albicans* and zymosan, revealed double-labeled cells, indicating phagocytosis of both types of particles (Fig. 3B, Supp. Fig. S13). Upon UV-inactivation, *C. albicans* retains its fungal cell wall intact, as in intestinal colonizing cells (Esteban et al., 2011; Galan-Diez et al., 2010). Uptake of labeled UV-inactivated *C. albicans* (of a different wild type strain-SC5314) by ileal and colonic organoids was observed, suggesting their capability to phagocytose luminal fungi (Fig. 3C; Supp. Fig S12B). Finally, Dectin-1 localization around intracellular HK-*C. albicans* (Fig. 3D), which is supported by intensity fluorescence profile (Fig. 3E), infers its involvement in phagocytosis. We cannot exclude the possibility that other phagocytic receptors may contribute to epithelial phagocytosis of *C. albicans*, hence, we asked whether such receptors are expressed by IECs. To this end, we tested Dectin-2/CLEC6A, which is implicated in fungal phagocytosis (Ifrim et al., 2016; Kitai et al., 2021; Lamprinaki et al., 2017). Indeed, we detected Dectin-2 on the surface of human ileal and colonic IECs isolated from surgical specimens by flow cytometry (Supp. Fig. S15 A-C), and in colonic frozen sections (Supp. Fig. S15 D) as well. This finding further supports the idea of epithelial capability to phagocytose luminal fungi although the role of Dectin-2 in phagocytosis was not addressed here.

**Figure 3:**
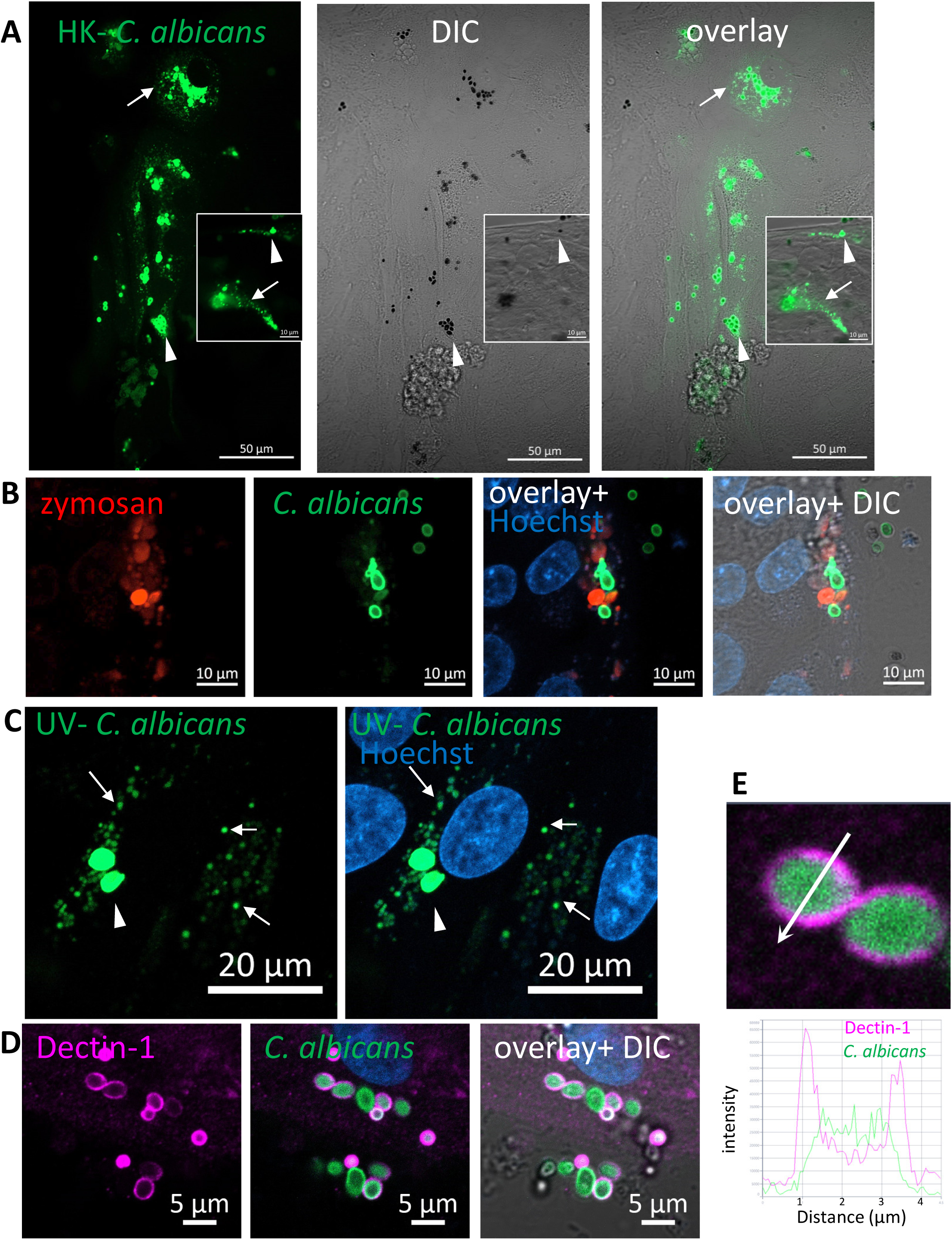
Phagocytosis of *C. albicans* by human intestinal organoids. Colonic organoids were fed over night with Rhodamine-green-X labeled HK-*C. albicans* (A, green), or both pHrodo-red zymosan and HK-C. albicans (B) or UV-inactivated *C. albicans* (C). Live confocal images were acquired directly or after nuclear stain with Hoechst 33342 (blue). Arrowhead - intact *C. albicans*, arrow-fragmented C. albicans. Original magnification x40, scale bars 50 µm (A), 10 µm (A-inset and B) and 20 µm (C). (D) Dectin-1 is recruited to phagocytosed *C. albicans*. Ileal organoids were fed with Rhodamine-green-X labeled HK-*C. albicans*. Following fixation organoids were stained with Dectin-1 polyclonal antibody. Original magnification x63, scale bar 5 µm. (E) Fluorescence intensity profile along the arrow of an inset from (D) is shown on the graph.

### LC3 is recruited to phagosomes of fungal particles

LC3 associated phagocytosis (LAP) is a receptor-mediated phagocytosis that utilizes some components of the autophagy machinery to process extracellular cargo (Grijmans et al., 2022; Heckmann and Green, 2019; Sanjuan et al., 2007). Dectin-1 mediated LAP has been demonstrated in phagocytosis of fungi by professional phagocytes such as macrophages and dendritic cells (Ma et al., 2012; Tam et al., 2014). To determine whether LAP of fungal particles occurs in IECs, we generated SW480 cells stably expressing GFP-tagged-LC3. These cells were incubated with pHrodo-red zymosan and using live confocal imaging pHrodo-red zymosan (Fig. 4A) engulfed by GFP-LC3 was detected, and was further confirmed upon analysis of fluorescence intensity profile (Fig. 4B). Similarly, using an antibody specific to LC3 LAP-engaged phagosomes (LAPosomes, (Martinez et al., 2015)) of zymosan and HK-*C. albicans* in organoids (Fig. 4C-D, Supp. Fig. S16A) and IEC lines were identified (Supp. Fig. S16B). Interestingly, the commercial preparate of HK-*C. albicans* contained a minor fraction of hyphae. We found that IECs could phagocytose and recruit LC3 to the hyphal form of *C. albicans* (Fig. 4E). This surprising finding suggests that IECs can recognize and internalize a wide range of microbial forms.

**Figure 4:**
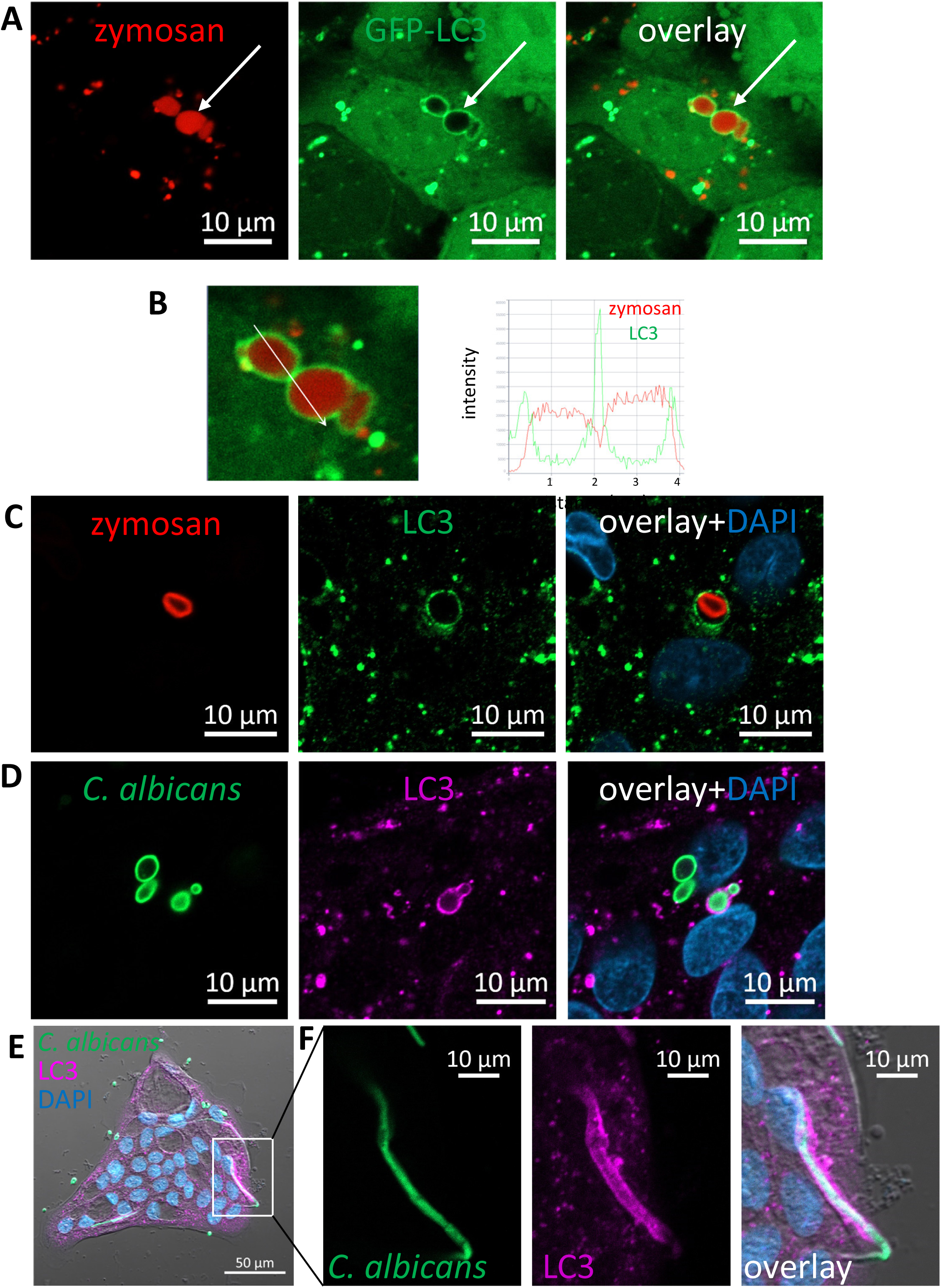
LC3 is recruited to phagosomes in IECs. (A) SW480 LC3-GFP cells were fed with pHrodo-red zymosan over-night. Live imaging shows LAPosomes (arrow) as LC3 (green) around intact zymosan (red) particles, as well as fragmented zymosan and autophagosomes. (B) Fluorescence intensity profile along the arrow of an inset from (A) is shown on the graph. (B) Colonic organoids were fed with pHrodo-red zymosan (red) over-night and stained with LC3 antibody (green) and DAPI (blue). (D-F) Colonic organoids were fed with Rhodamine-green-X HK-*C. albicans* (green) over-night and stained with LC3 antibody (magenta) and DAPI (blue). Shown is LAP of yeast (D) and hyphal form (E,F) of HK-*C. albicans*. F is an inset of E. Original magnification ×40 (A, B), ×63 (C,D) x20 E,F) scale bar 10 µm (A-D) and 50 µm (E).

### Rubicon is recruited to phagocytosed zymosan and *C. albicans*

Rubicon is a key regulatory protein considered unique to LAP in professional phagocytes (Boyle and Randow, 2015; Wong et al., 2018). In order to support the notion that IECs are capable of LAP we assessed the presence of Rubicon at zymosan and *C. albicans*’ particles upon their incubation with organoids. Indeed, using a specific antibody we identified Rubicon around intracellular HK-*C. albicans* (Fig. 5A-B) and zymosan (Fig. 5C-E) in intestinal organoids, suggesting its recruitment and involvement in the phagocytic process, lending further support to the occurrence of LAP in IECs.

**Figure 5:**
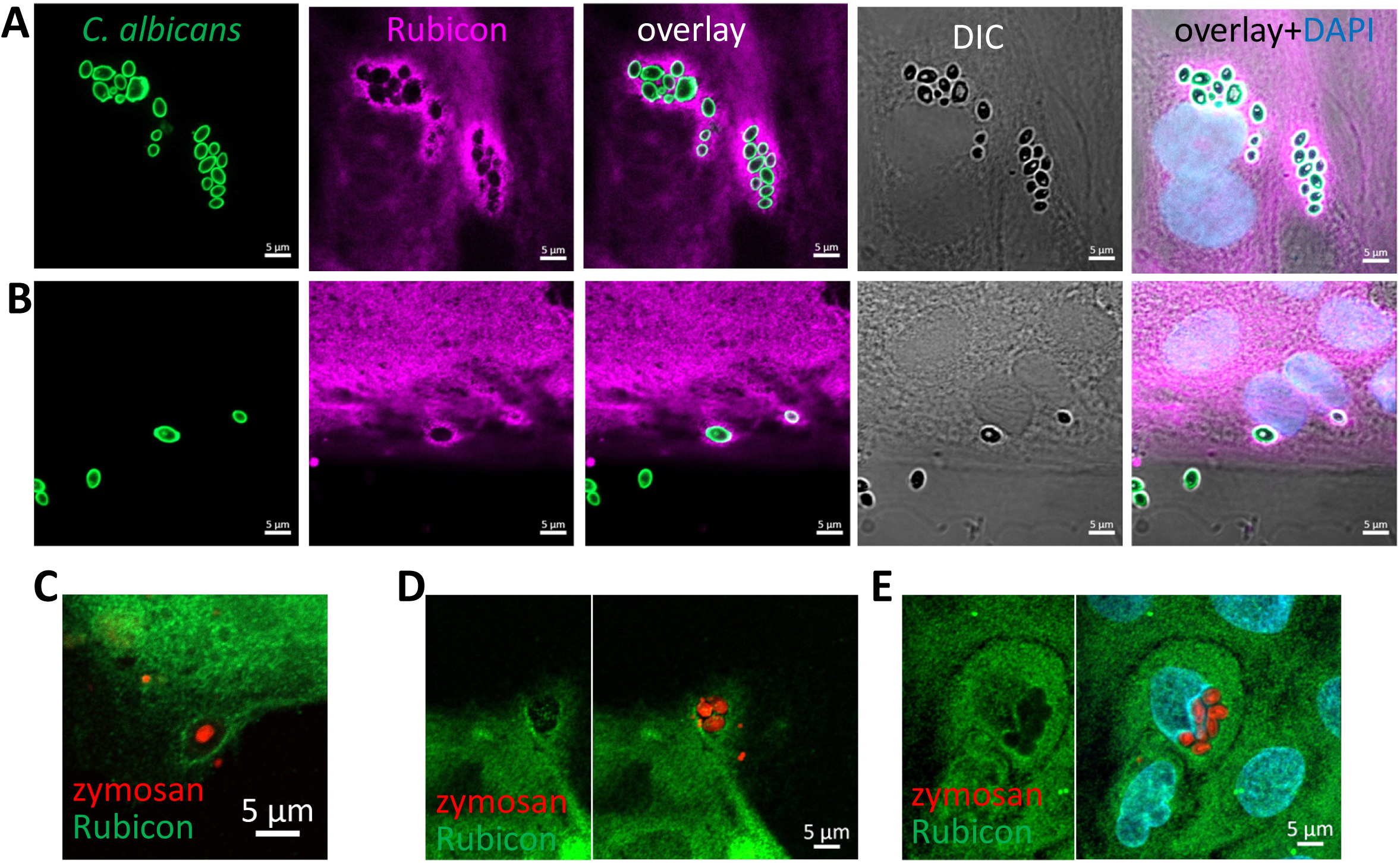
Rubicon is recruited to the phagosome. Colonic (A) and ileal (B) organoids from the same individual were fed over-night with HK-*C. albicans* (green) and stained with Rubicon antibody (magenta) and DAPI (blue). (C-E) Colonic (C) and Ileal (D-E) organoids were fed with pHrodo-red zymosan (red) and stained with Rubicon (green) and DAPI (cyan). Original magnification x63, scale bar 5 µm.

### Phagocytosis depends on NADPH-oxidase activity

A hallmark of LAP in macrophages is the production of reactive oxygen species (ROS) by nicotinamide adenine dinucleotide phosphate (NADPH) oxidase-2 (NOX2) (Heckmann and Green, 2019; Martinez et al., 2015). Human colonic IECs express NOX1, which is a structural homolog of NOX2 as well as additional NADPH oxidases including DUOX2 (Schwerd et al., 2018). To test whether NADPH-oxidases are involved in IEC-mediated phagocytosis, we treated organoid monolayers and SW480 cells with pHrodo-red zymosan after exposure to the general NADPH oxidase inhibitor diphenyleneiodonium (DPI). Figure 6A demonstrates that DPI (at 2 µM) drastically suppressed zymosan phagocytosis in organoids. This finding is supported by quantification of phagocytosis as reflected by total fluorescence using a microplate-reader where up to 44% and 65% inhibition by 2 µM (***p≤ 0.001) and 10 µM DPI (****p≤ 0.0001) respectively was observed (Fig. 6B) and similar inhibition was observed cell lines (Fig. 6C). Our results indicate that NOX activity is necessary for phagocytosis, implying a role for ROS production and supports the occurrence of LAP in IECs.

**Figure 6:**
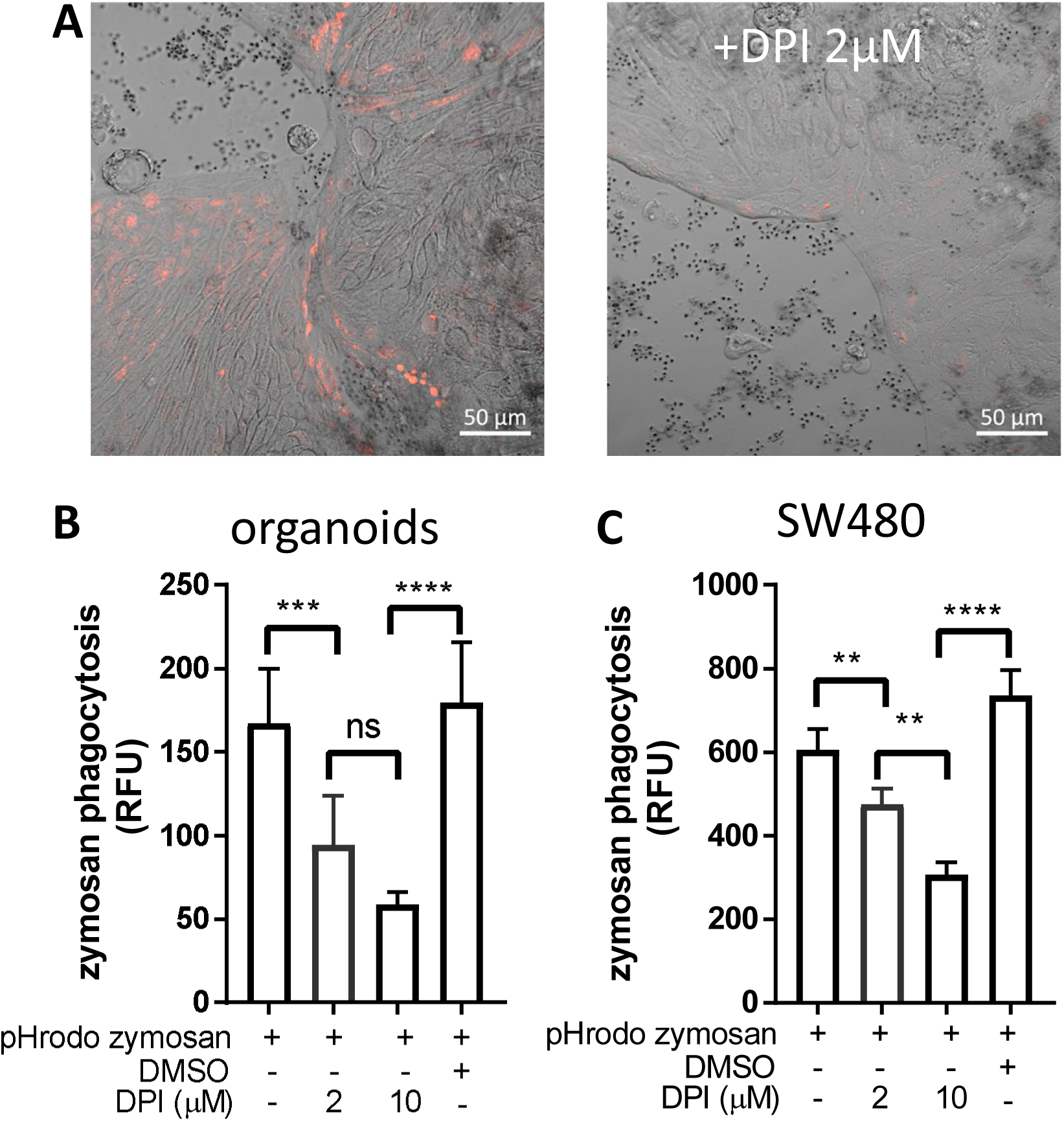
DPI inhibits zymosan phagocytosis. (A) Colonic organoid monolayers grown in differentiation medium for 3 days were exposed to pHrodo-red zymosan for 24 hours in the presence or absence DPI (2 µM) that was added for 60 minutes prior to the addition zymosan. Shown are intracellular fluorescent zymosan (red) and the organoid cells (DIC) of representative frames from 7-15 random fields imaged. The experiment was repeated 3 times using 2 different organoids. Original magnification x20, scale bar 50 µm. (B) Colonic organoids were seeded in 96 well plate, treated as in (A) or in the presence of vehicle (DMSO 1:1000) in 8-replicate wells for 48 hours. Phagocytosis was assessed as the relative fluorescence by a microplate reader. (C) SW480 cells were seeded in 96 well plate and treated as in (B) in triplicates for 24 hours. Data are shown as mean ± SD of biological 8-replicates (B) and triplicates (C) from a representative of three independent experiments performed. **p<0.01, ***p<0.001, ***p<0.0001, One-way ANOVA followed by Tukey multiple comparison test.

### Phagocytosed fungi are degraded in the lysosomes

The evidence regarding intracellular fragmentation of zymosan and *C. albicans* (Figs. 1-3), suggest the occurrence in of phagosome maturation via fusion with lysosomes and lysosomal degradation of phagosome content. To verify this, we asked whether phagocytosed zymosan and *C. albicans* colocalize with lysosomes. Indeed, intact and fragmented zymosan and *C. albicans* (both HK- and UV-inactivated) colocalized with acidic organelles as identified by lysosomal dyes in ileal and colonic organoids and in cell lines (Fig. 7A-C, Supp. Fig. S17A -B). In addition, the lysosomal protein LAMP2 encircled phagocytosed particles in organoids and cell lines (Fig. 7D-E, Supp. Fig S17C). Finally, colocalization of LAMP2 with LC3 around intracellular zymosan indicates LAPosomes that fuse with lysosomes (Fig. 7F-G).

**Figure 7:**
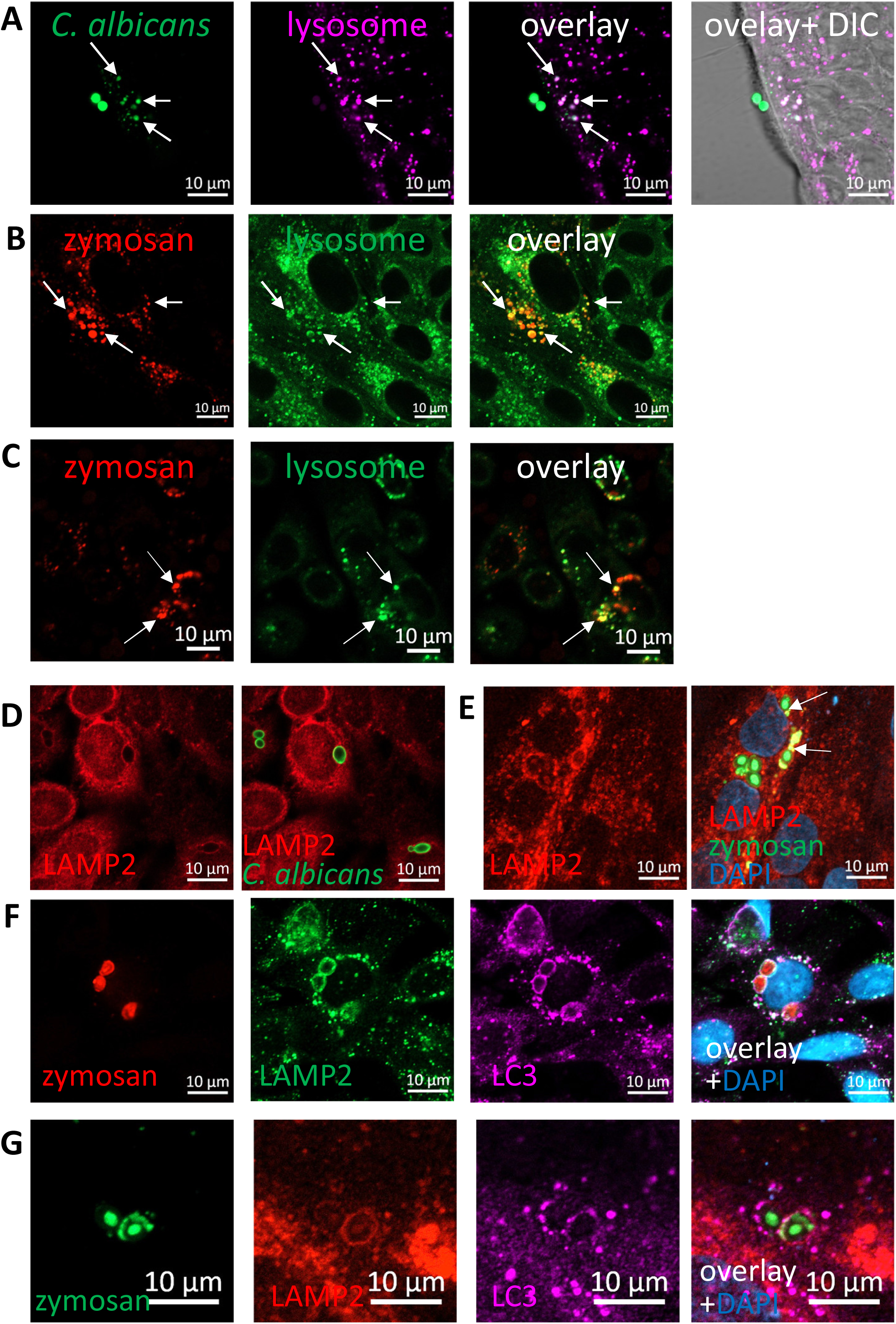
Phagocytosed particles are directed to lysosomal processing. (A) Colonic organoids were incubated over-night with Rhodamine-Green labeled HK-*C. albicans* (green) and stained with lysosomal-NIR reagent (magenta). (B, C) Ileal organoids (B) and SW480 cells (C) were incubated with pHrodo-red zymosan (red) and stained with lysosomal-green reagent. Arrows indicate colocalization of fragmented HK-*C. albicans* or zymosan and lysosomes. (D) SW480 cells were fed with HK-*C. albicans* (green) and stained with LAMP2 antibody. (E) Intact and fragmented zymosan particles are surrounded by LAMP2. Ileal organoids were fed with AF488-zymosan (green) over night, and stained with LAMP2 antibody (red). Arrowhead - intact zymosan, arrow-fragmented zymosan. (F, G) LAPosomes merge with lysosomes. (F) SW480 cells were fed with pHrodo red zymosan (red) and stained with LAMP2 (green) and LC3 (magenta) antibodies and counterstained with DAPI (blue). (G) Ileal organoids were fed with AF488-zymosan (green) and stained with LAMP2 (red) and LC3 (magenta) antibodies and counterstained with DAPI (blue). Original magnification x63 (A,D-G), x40 (B), x20 (C), scale bar 10 µm.

Together, our findings suggest that human IECs are capable of LAP of fungal particles, and provide mechanistic evidence for stages of the process, starting from their identification by the receptor, Dectin-1, via recruitment of Rubicon and LC3 to final degradation in the lysosomes.

## Discussion

The intestinal epithelium acts as a physical and functional barrier as well as an active participant in mucosal immunity by orchestrating protection against pathogens and maintaining tissue homeostasis (Gunther and Seyfert, 2018; Soderholm and Pedicord, 2019). We and others had previously shown that IECs exert various responses upon interaction with bacterial and fungal components through pattern-recognition receptors such as Toll-like receptors (TLRs) and Dectin-1, yet they appear to tolerate various commensal microorganisms (Burgueño and Abreu, 2020; Cohen-Kedar et al., 2014; Cohen-Kedar et al., 2021; Ladinsky et al., 2019; Richardson et al., 2018).

In this work we present a novel mode of IEC-microbiota interaction where IECs along the human intestine can internalize commensal fungal particles via LAP. Importantly, our data demonstrate that this is a host-driven process, since the phagocytosed particles are fully inactive. As non-professional phagocytes, the phagocytic capability of epithelial cells is considered limited compared to professional phagocytes (Freeman and Grinstein, 2016; Gunther and Seyfert, 2018). However, their abundance on large mucosal surfaces may contribute to tissue homeostasis and the defense against pathogens (Gunther and Seyfert, 2018). Indeed, previous studies in human and murine models demonstrated that epithelial cells (colonic, mammary and hair follicular) engulf apoptotic cells (Fornetti et al., 2016; Lee et al., 2016; Mesa et al., 2015). Moreover, Retinal pigment epithelium used LAP to clear photoreceptor outer segments in mice (Kim et al., 2013; Muniz-Feliciano et al., 2017) and HeLa cells used a host derived LAP-like mechanism to target *Yersinia Pseudotuberculosis* (Ligeon et al., 2014).

In this work we focused on fungal recognition by IECs predominantly via the interaction of Dectin-1 with fungal β-glucan. However, additional phagocytic receptors such as Dectin-2 (this report) and TLRs (Burgueño and Abreu, 2020) are expressed by IECs and a previous work further demonstrated TLR4 mediated bacterial phagocytosis by mouse enterocytes (Neal et al., 2006). Together these findings support the notion of epithelial capability to phagocytose luminal microorganisms.

LAP of fungi and fungal materials has been characterized in professional phagocytes (Cadwell, 2016; Oikonomou et al., 2018; Sprenkeler et al., 2016) such as mouse macrophages and dendritic cells (Ma et al., 2012; Tam et al., 2014) and in human monocytes (Kyrmizi et al., 2013). The role of Dectin-1 in this process was demonstrated by the inhibition of LAP upon Dectin-1 deficiency or following exposure to its antagonist, laminarin (Kyrmizi et al., 2013; Ma et al., 2012). Likewise, we demonstrate in this work that Dectin-1 was involved in zymosan phagocytosis by human IECs. While Syk is activated by zymosan and fungi in IECs (Cohen-Kedar et al., 2014; Cohen-Kedar et al., 2021; Wang et al., 2021) it seems dispensable for phagocytosis. Examples for Dectin-1 mediated Syk-independent phagocytosis exist also in macrophages (Herre et al., 2004; Underhill et al., 2005).

Recruitment of LC3 to the internalized zymosan and *C. albicans* yeast and hyphal forms is indicative for LAP. Furthermore, Rubicon, that acts as a regulatory switch to inhibit autophagy and to promote LAP (Boyle and Randow, 2015; Wong et al., 2018), engulfs intracellular zymosan and *C. albicans* suggesting its active role in IECs’ phagocytosis. Finally, the sensitivity of phagocytosis to inhibition of NADPH-oxidases denotes a role for ROS production. In macrophages, the phagocytic NOX2 complex plays a role in LAP, but not in canonical autophagy (Heckmann and Green, 2019).

While NOX2 is not expressed by IECs, it might be replaced by other NADPH-oxidases, such as NOX1. Together these findings show that phagocytosis is a regulated physiological process in IECs. There is scarce evidence for fungal phagocytosis by epithelial cells: Dectin-1 mediated phagocytosis of spores of *Aspergillus fumigatus* has been reported in airway epithelium (Bertuzzi et al., 2014), and zymosan was internalized by chicken IECs in 3D organoids (Nash et al., 2021). Yet, neither LAP nor degradation of commensals have been previously shown in primary human epithelial cells.

Luminal sampling at steady state is important for homeostasis and building mucosal tolerance towards commensal microorganisms and is usually attributed to professional mucosal-resident phagocytes and to transcytosis (uptake and delivery without intracellular degradation) by specific epithelial M-cells, that expose immune cells in the lamina propria to luminal antigens (Chu et al., 2016; Grijmans et al., 2022; Rios et al., 2016; Schulz and Pabst, 2013; Yu et al., 2014). The phagocytic capacity presented in this work is not likely to be assigned to M-cells for several reasons: first, it is observed in three human epithelial cell lines and in organoids derived from different parts of the gastro-intestinal tract from ileum to sigmoid colon, while M-cells are mostly found in the small intestine and require specific differentiation conditions including TNFα and RANKL (Fasciano et al., 2019; Hsu et al., 2022; Staab et al., 2020) that were not used in this study. Second, we find that phagocytosis occurs by MUC2 positive and MUC2 negative cells indicating goblet and non-goblet cells. Finally, we find evidence for Dectin-1 mediated LAP, that involves intracellular epithelial processing of the internalized particle, as opposed to the transcytosis of intact particles by M-cells.

The role of commensal fungi in shaping mucosal tolerance and host systemic immune response has been recently established in a series of reports (Doron et al., 2021a; Doron et al., 2021b; Leonardi et al., 2022; Li et al., 2022) underlying their interaction with mucosal immune cells e.g., lymphocytes via mononuclear phagocytes. Here we propose epithelial phagocytosis of luminal fungi as another pathway for intestinal mucosal sensing of fungi. While the physiological outcome of epithelial phagocytosis and lysosomal degradation of fungi is still uncovered, it is plausible to assume that it has an impact on the mucosal milieu. Indeed, an example where IECs acquire antigens from commensal bacteria, (segmented filamentous bacteria, SFB), for generation of TH17 cell responses in mice was recently provided (Ladinsky et al., 2019). IECs did not phagocytose SFB, nor did SFB penetrate their cytosol. Rather, they endocytose vesicles containing SFB cell wall–associated proteins, that acted as immunomodulators on T cells. Still, the mechanism by which IECs induced Th17 differentiation remains unclear. Interestingly, mucosa-associated fungi elicit a protective effect on IECs barrier function and protected mice against intestinal injury and bacterial infection via Th17 cells (Leonardi et al., 2022). It is not known whether IECs play an active role in the observed increased frequency of Th17 cells.

In professional phagocytes, LAP of fungi enhances pathogen killing (Tam et al., 2014) (Kyrmizi et al., 2013), and cytokine secretion (Lamprinaki et al., 2017). Also, it promotes MHC class II recruitment to the LAPosome leading to sustained antigen presentation (Ma et al., 2012).

IECs express MHC class II and co-stimulatory molecules and hence were suggested to act as antigen presenting cells (APCs) (Beyaz et al., 2021; Hershberg and Mayer, 2000; Heuberger et al., 2021; Rabinowitz and Mayer, 2012). Indeed, there is evidence that MHC class II-expressing IECs functioned as APCs to prime donor CD4+ T cells *ex-vivo* and *in vivo* where microbiota influences MHC class II expression on IECs in the ileum. Still, the presented peptides were mostly thought as endogenous peptides (Koyama et al., 2019). Whether IECs can present exogenous peptide is not clear, and an experimental model was presented to study the interactions between IEC MHC-II and the surrounding immune and microbial milieu (Wosen et al., 2019). We propose that degradation products of phagocytosed fungi might be presented in context with MHC class II by IECs, similarly to their presentation upon phagocytosis by professional antigen presenting cells. If IECs indeed present fungal antigens, they may contribute to humoral responses. It was recently found that commensal fungi induce, via intestinal mononuclear phagocytes, the production of secretory IgA (sIgA) in the murine gut and systemic serum IgG (Doron et al., 2021a; Doron et al., 2021b). A decrease in antifungal sIgA was observed in patients with Crohn’s disease (Doron 2021), underlying a feature of loss of tolerance towards commensal microorganisms, which is typical of inflammatory bowel diseases (IBD) (Iliev and Cadwell, 2021).

Accordingly, possible contribution of interrupted epithelial LAP to disrupted mucosal homeostasis and loss of tolerance may be assumed. An attractive approach will be to assess whether genes associated with IBD that may function in LAP indeed affect epithelial LAP. An immediate candidate from our work is NOX1, which has recently been shown to prevent inflammation, and its mutations were linked to ulcerative colitis (Hsu et al., 2022; Schwerd et al., 2018). Another interesting candidate is the ATG16L1 T300A polymorphism. Individuals carrying this polymorphism exhibit defects in T-regulatory responses to outer-membrane-vesicles (OMVs) of the commensal *Bacteroides fragilis*, and sensing those OMVs, in mouse dendritic cells, occurs via LAP and involves Rubicon and ATG16L1 (Chu et al., 2016).

Our findings may have mechanistic and translational implications, as well as paving the way for detailed characterization of the processes underlying epithelial phagocytosis of microorganisms. Specifically, facilitating further identification of host and microbial factors, such as cell wall composition of pathogens compared to commensals, that control epithelial phagocytosis. Our experimental setting may also be useful to study the outcome of IECs’ phagocytosis with respect to the impact on neighboring cells (e.g., T-cells). Using patient derived organoids (Fujii and Sato, 2021), our findings may allow the assessment of epithelial phagocytic capabilities under defined genetic (e.g. ATG16L1 variants) and clinical (IBD) states and evaluate their contribution to homeostasis or if perturbed, to the pathogenesis of IBD.

## Methods

### Cell lines

Human colon epithelial cell lines SW480, HCT116 and Caco-2 were purchased from ATCC (Manassas, VA). SW480 and HCT116 cells were grown in RPMI medium supplemented with 10% fetal bovine serum, and Caco2 cells were grown in EMEM, supplemented with 20% fetal bovine serum (from Biological Industries). All growth media contained 100 units/mL penicillin G, and 100 µg/mL streptomycin. All cells were maintained in a humidified incubator at 37°C with 5% CO2.

To generate cells stably expressing GFP-LC3, SW480 cells were transfected with pEGFP-LC3 using the Lipofectamine 2000 reagent (Invitrogen) according to the manufacturer’s instructions. Stable clones expressing GFP-LC3 were selected and cultured in the presence of 1000 μg/ml geneticin (G-418, Calbiochem). pEGFP-LC3 (human) was a kind gift from Toren Finkel (Lee et al., 2008) (Addgene plasmid # 24920).

### Human samples and ethics statement

The Institutional Ethical Committee of the Rabin Medical Center approved the study (approval number 0763-16-RMC and 0298-17) and a written informed consent of all participating subjects was obtained. The identity of all participating subjects remained anonymous. Tissue samples were taken from surgical specimens of patients undergoing bowel resection for colonic tumors (normal ileal or colonic samples were taken from a distance of at least 10 cm from the tumor). Specimens were kept over-night at 4°C in RPMI containing 100 units/mL penicillin G, and 100 µg/mL streptomycin and 2.5 µg/mL amphotericin B (Fungizone) supplemented with 10% fetal bovine serum.

### Human intestinal organoids

Human ileal and colonic crypts were isolated and organoids were cultured as previously described (Sato et al., 2011). In brief, tissue fragments were washed twice with crypt isolation medium: 0.5 mM DL-Dithiothreitol (DTT), 5.6 mM Na2HPO4, 8 mM KH2PO4, 96.2 mM NaCl, 1.6 mM KCl, 43.4 mM sucrose, 54.9 mM D-sorbitol.

Tissue was incubated in crypt isolation medium supplemented with 2 mM EDTA, for 30 minutes at 4°C followed by vigorously shaking till crypt were released from the mesenchyme. Crypt pellet was washed with FBS and resuspended in ice-cold Matrigel and seeded as 15 µl domes on pre-warmed 12-wells tissue culture plates. Plates were incubated upside down for 20 min in a 37°C 5% CO2 incubator until the Matrigel solidifies. Organoid expansion media (based on (Pleguezuelos-Manzano et al., 2020; Usui et al., 2018)) consisted of advanced DMEM F12 (Gibco) (26% of total volume), 100/100 U/ml Penicillin/streptomycin, 10 mM HEPES, 1× GlutaMAX, and the following growth factors: 1× B27, 1 mM N-Acetylcysteine, 100 ng/ml Noggin, 50 ng/ml human EGF, 10 mM Nicotinamide, 10 μM SB202190, 500 nM A83-01, 10 nM Prostaglandine E2, 26 μg/ml Primocin, and conditioned medium from the L cell line secreting Wnt3A (50% of total volume) and 293T cells secreting R-spondin 1 (20%). 10 μM Y-27632 (Rock inhibitor) was added to expansion medium for the first 2 days. Medium was changed every other day.

2D organoid monolayer culture protocol was based on the supplementary protocol of Intesticult™ medium (WWW.STEMCELL.COM) culture: µ-Slide 8-well glass bottom chambers (Ibidi, Martinsried, Germany) were coated with 1:50 Matrigel in PBS for one hour at a 37°C 5% CO2 incubator. 3D organoids were mechanically disrupted into a single cell suspension. Cells were resuspended in expansion medium containing 10 μM Y-27632 and seeded on coated slides (approximately 2 domes per 8-well chamber). Expansion medium was changed every other day until 2D organoids reached 50-70% confluency. Prior to phagocytosis experiments, 2D organoids were grown for additional 2-3 days in a generic differentiation medium based on (Pleguezuelos-Manzano et al., 2020). Briefly, advanced DMEM F12 was supplemented with 100/100 U/ml Penicillin/streptomycin, 10 mM HEPES, 1× GlutaMAX, 1× B27, 1 mM N-Acetylcysteine, 500 ng/ml human R-spondin 1, 100 ng/ml Noggin, 50 ng/ml human EGF, and 100 μg/ml Primocin.

### *Candida albicans* growth conditions and labeling

*C. albicans* wild type strain SC5314 was kindly provided by Judith Berman (Tel Aviv University). Cells were grown over-night in YPAD medium at 30°C. UV inactivated *C. albicans* cells were prepared as previously described (Wheeler and Fink, 2006). Briefly, cells were exposed in a thin liquid suspension to 4 doses of UV radiation (100 mJ/cm^2^) in a UV cross linker (CL-1000 UVP, Upland, CA). Cells were counted and resuspended in PBS. Killing was verified by seeding onto YPAD-agar plates. Heat-killed *C. albicans* wild type strain ATCC 10231 (tlrl-hkca) was purchased from InvivoGen (San Diego, CA) and resuspended in endotoxin-free water.

Heat-killed or UV-inactivated particles of *C. albicans* were labeled with Rhodamine Green-X (Life Technologies, Invitrogen, R-6113) as previously described (Marakalala, 2014). After labeling cells were washed, resuspended in PBS, counted and aliquoted.

### Confocal phagocytosis assay

IEC lines were seeded on μ 8-well glass-bottomed slides (Ibidi, Martinsried) at a density of 3-4x10^4^ cells/well. Two days later, medium was replaced and cells were stimulated over-night with pHrodo-red zymosan (100 µg/ml, P35364 Invitrogen) or AF488-zymosan (100 µg/ml) Z23373 Invitrogen) or Rhodmine-green-x labeled *C. albicans* (approximately at 0.3-1×10^6^/well). We counted about 10^5^ cells/well, hence, MOI= 3-10 /cell). The next day cells were either directly imaged with confocal microscope (LSM 800, Zeiss, and Zen3.2 software) or stained with Hoechst 33342 (14533, Sigma) to visualize nuclei or with lysosomal staining reagents (Green - Cytopainter ab176826 or NIR - Cytopainter ab176824, Abcam). For image analysis and fluorescence intensity profile we used Zen 3.2 software. Where indicated, the inhibitors Cytochalasin-D (10 µM, MBS250255, Calbiochem) or laminarin (1 mg/ml, L9634, Sigma) or diphenyleneiodonium (DPI) (2 µM, D2926, Sigma) were added 1 hour prior to pHrodo-red zymosan addition.

2D organoid monolayers were seeded as described above, and grown in differentiation medium for at least 2 days prior to their exposure to labeled zymosan or *C. albicans.* Then, organoids were treated and assessed as the cell lines. In some cases, cells or organoids were fixed with 4% paraformaldehyde or 100% methanol for further immuno-staining. All images within each experiment were acquired under the same conditions.

### Image quantification of phagocytosis

We quantified phagocytosis either by confocal image analysis using ImagJ or by multi-well-plate fluorescence quantification using a microplate reader. Image-analysis quantification was applied on confocal images of cell lines, where random fields acquired were representative of the uniformly distributed phagocytic cells. For each experimental condition (phagocytosis in the presence or absence of inhibitors) the number of nuclei within at least four confocal random fields (acquired at x20 magnification) was determined with ImageJ. Then the number of pHrodo-red zymosan positive cells within each field was manually counted. The percentage of positive cells was calculated. For each condition, at least 2000 cells were analyzed.

### Microplate reader quantification of phagocytosis in cell lines and organoids

We assessed the level of phagocytosis by quantification of the total fluorescence within replicate wells as an additional assay for cell lines, and as the main assay quantify phagocytosis in organoid monolayers where it was highly important to quantify the whole well since phagocytosis is not uniformly distributed. SW480 and HCT116 cells were seeded in 96-well flat-bottomed plastic plates at a density of 4x10^4^ cells/well for 24 hours. The next day the medium was replaced with 100 µl/well of fresh medium or medium that contains inhibitors (Cytochalasin-D 10 µM, Laminarin 1 mg/ml, Syk inhibitor 574711 [3-(1-methyl-1H-indol-3-yl-methylene)-2-oxo-2,3-dihydro-1H-indole-5-sulfonamide], (Calbiochem, Merck-Millipore) 1 or 5 µM, DPI (2 or 10 µM) or DMSO as vehicle (1:1000) where applicable and pHrodo-red zymosan at 100 µg/ml was added in triplicate wells for 24 hours. Wells were washed gently 3 times with white-RPMI and relative fluorescence was measured with Synergy H1 microplate reader (Biotek). For phagocytosis quantification in organoids, 96-well plastic plates were coated with matrigel as described for 8-well chambers.

Cells from 3D human colonic organoids were resuspended and seeded in a ratio of one-two domes to 60 wells or 10,000 cells/well. Organoid monolayers were grown in expansion medium for at least 3 days, reaching at least 50% confluence, and then the medium was replaced with differentiation medium for 2-3 days (where wells were 80-90% confluent). Before stimulation, medium was replaced with fresh differentiation medium in the presence or absence of laminarin (1 mg/ml) or DPI (2 or 10 µM), and pHrodo-red zymosan at 100 µg/ml was added in triplicates to 8-replicate wells as indicated for 48 hours to allow accumulation of phagocytic cells thus obtaining enhanced phagocytosis signal. Medium was removed and organoids were washed gently 3 times with white-RPMI before relative fluorescence was assessed in a microplate reader.

### Cytokine secretion (ELISA)

SW480 cells were seeded in 96-well flat-bottomed plastic plates at 4x10^4^ cells/well for 24 hours. The next day the medium was replaced with fresh medium or medium that contains Syk inhibitor (1 or 5 µM/ml) (100 µl/well) and one hour later, zymosan (100 µg/ml, tlrl-zyn, InvivoGen, not labeled) was added in triplicate wells for 24 hours. Supernatants were assessed for IL-8 secretion using ELISA (DY208 R&D systems) according to the manufacturer’s instructions. In parallel, in the same 96-well plate, triplicate wells were exposed to pHrodo-red zymosan in the presence and absence of the same Syk inhibitor, for phagocytosis microplate reader quantification assay.

### Cell staining by Immunofluorescence

Cells or organoid monolayers seeded in 8-well chambers and treated as indicated in the figure legends, were fixed with 4% paraformaldehyde for 30 minutes at room temperature or with ice-cold methanol for 15 minutes at -20°C according to the requirement of the primary antibodies, washed with PBS, and blocked-permeabilized with 5% donkey serum containing 0.3% triton for 1 hour at room temperature. Cells or organoid monolayers were incubated with the indicated primary antibodies overnight at 4°C, followed by staining with the corresponding fluorescently labeled secondary antibodies for 1 hour at room temperature and counterstaining with DAPI, which was included in the non-hardening mounting medium (GBI Labs, E-19-18, Mukilteo, WA). Samples were visualized by inverted confocal microscope (LSM 800, Zeiss). All images within each experiment were acquired under the same conditions. Details of primary and secondary antibodies appear Table 1.

**Table 1:** Antibodies used for immuno-fluorescence

| Antibody | host | Fixation | Dilution | Cat. # | Company |
| --- | --- | --- | --- | --- | --- |
| Dectin-1 | Rabbit polyclonal | Methanol | 1:200 | NBP1–25514 | Novus Biologicals |
| Dectin-1 (GE2) | Mouse monoclonal | PFA | 1:50 | ab82888 | Abcam |
| LC3A/B (D3U4C) | Rabbit monoclonal | Methanol | 1:100 | 12741 | Cell Signaling |
| Rubicon | Mouse monoclonal | PFA | 1:100 | ab156052 | Abcam |
| LAMP2 (H4B4) | Mouse monoclonal | Methanol | 1:200 | sc-18822 | Santa cruz |
| Ki67 | Rabbit monoclonal | PFA | 1:400 | RBK027 | Zytomed |
| Muc2 (996/1) | Mouse monoclonal | PFA | 1:200 | ab11197 | Abcam |
| Lysozyme | Rabbit polyclonal | PFA | 1:400 | A 0099 | Dako |
| Dectin-2 | Mouse monoclonal | PFA | 1:100 | ab107572 | Abcam |
| ZO-1/TJP1 | Mouse monoclonal | PFA | 1:100 | 33-9100 | Thermo Scientific |
| EpCAM (VU-1D9) | Mouse monoclonal | PFA or Methanol | 1:400 | 187372 | Abcam |
| EpCAM | Rabbit polyclonal | PFA or Methanol | 1:400 | ab71916 | Abcam |
| anti mouse AF488 | Donkey |  | 1:1000 | ab150109 | Abcam |
| anti mouse AF647 | Donkey |  | 1:1000 | ab216773 | Abcam |
| anti rabbit AF488 | Donkey |  | 1:1000 | ab150065 | Abcam |
| anti rabbit AF647 | Donkey |  | 1:1000 | ab150067 | Abcam |

### Statistical analysis

Data are reported as the mean of triplicates ± SD of a representative example of at least 3 experiments. Significance was determined using unpaired tow-tailed student’s *t*-test or one-step ANOVA as indicated (GraphPad Prism 7.03 software, San Diego, CA). Differences were noted as significant by the following conventions: *p<0.05; **p<0.01; ***p<0.001 ****p<0.0001, as specifically indicated in the legend of each figure.

## Acknowledgments

The Authors would like to deeply thank Rabin Medical Center Institutional Biobank and for their invaluable support of this research. We thank Judith Berman from Tel Aviv University for *C. albicans* strains. This work was partially supported by a generous grant from The Leona M. and Harry B. Helmsley Charitable Trust to I.D. (Grant Number #2019PG-CD024).

## Supplementary Material

### Supplementary Methods

#### Fluorescent immunohistochemistry of frozen sections

Mucosal samples were taken from surgical specimens of patients undergoing bowel resection for colonic tumors. Normal mucosa was taken from a distance of at least 10 cm from the tumor and was snap-frozen in Tissue–Plus OCT compound (Scigen scientific Gardena) and kept at -80°C until sectioning for immunofluorescence.

Frozen sections (10 µM) were stained using a standard protocol according to the procedure in “Immunofluorescence” in the main text.

#### Isolation of primary human IECs for flow cytometry

IECs were freshly isolated as previously described (Brimnes et al., 2005; Rimoldi et al., 2005) with modifications. Briefly, mucosa and submucosa were separated. The tissue was dissected and incubated in 30 mL of HBSS without Ca2+ and Mg2+, containing 1 mM dithiothreitol, 5% FBS and 2 mM EDTA for 30 min at 37°C, while agitating at 250 rpm. The suspension was filtered through nylon mesh (70 mm pore diameter) and the pellet containing IECs was used flow cytometry.

#### Flow cytometry

Cell surface immunostaining of IECs was performed using APC-conjugated anti-human Dectin-2 mAb (clone 545943, FAB3114, R&D systems) according to the manufacturer’s instructions. All studies were carried out using matched isotype controls. Primary IECs were identified by co-staining with APC-conjugated human EpCAM mAb (clone 9C4, 324208, Biolegend). Data acquisition was performed using a FACSCanto flow cytometer (Becton Dickinson) and the data were analyzed with FlowJo software (Treestar, Inc.).

## Supplementary figure legends

**Supp. Fig. S1:**
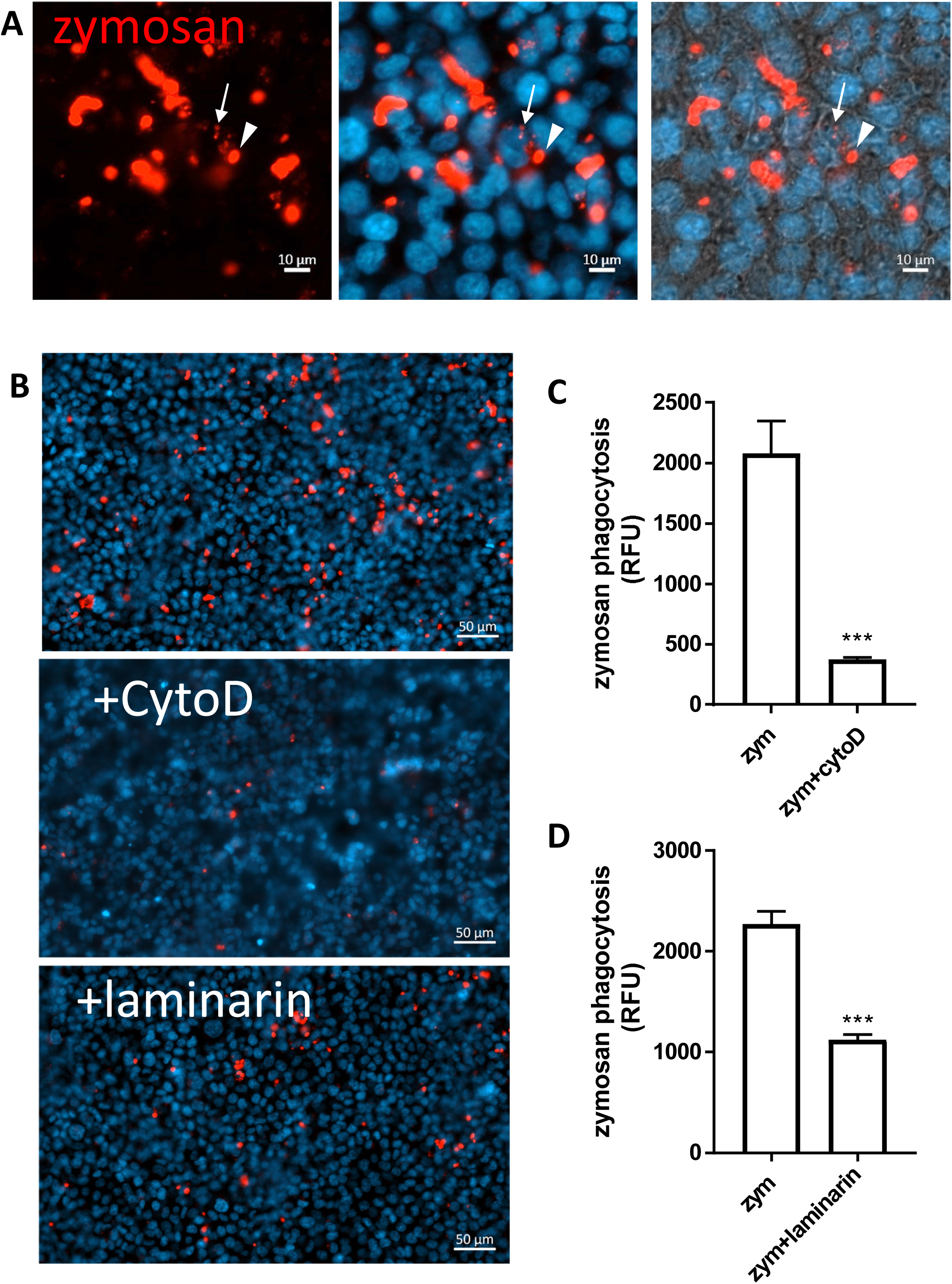
HCT116 cells uptake of zymosan depends on actin polymerization and Dectin-1. (A) HCT116 were fed with pHrodo-red zymosan over-night and stained with Hoechst 33342 prior to live imaging. Arrowhead – intracellular fluorescent zymosan, arrow-intracellular fragmented zymosan. Original magnification x20, scale bar 10 µm. (B) HCT116 were fed with pHrodo-red zymosan over-night in the presence or absence of Cytochalasin D (10 µM) or laminarin (1 mg/ml) which were added 1 hour prior to zymosan, and stained with Hoechst 33342 before live imaging. Original magnification x20, scale bar 50 µm. (C,D) HCT116 cells were seeded in 96 well plate and treated as in (B) and phagocytosis was assessed as the relative fluorescence by a microplate reader. Data are shown as mean ± SD of biological triplicates from a representative of three independent experiments performed.

**Supp Fig. S2:**
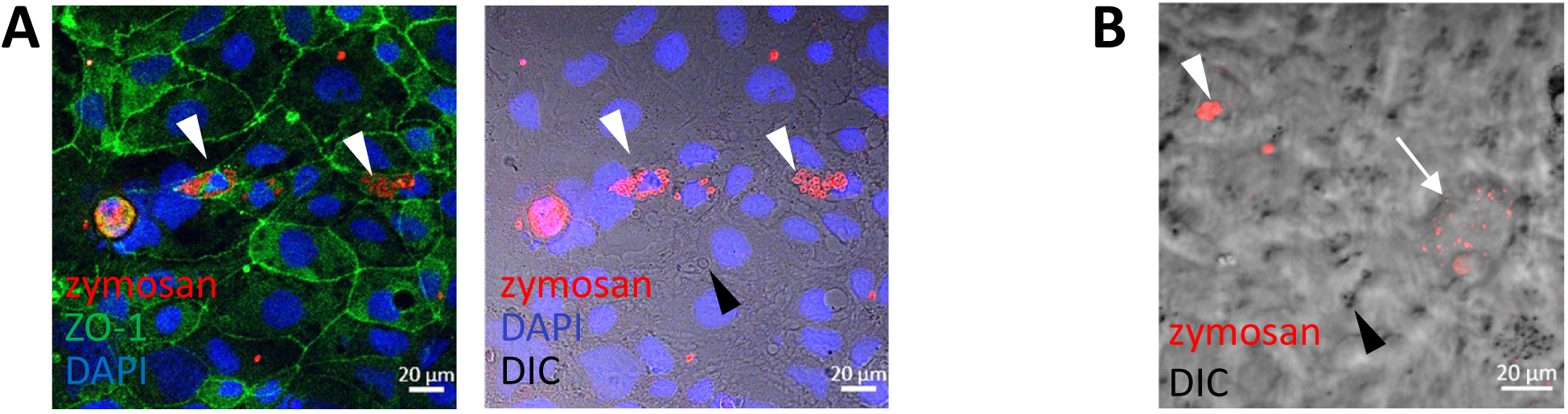
Caco-2 cells internalize and process zymosan. (A) Caco-2 cells were seeded on glass slides and fed with pHrodo-red zymosan over-night. Following fixation cells were immuno-stained with zo-1 to mark cell borderline (green) and nuclei were counterstained with DAPI (blue). Multiple intracellular zymosan particles are shown (white arrowheads) as well as extracellular zymosan (black arrowhead). Original magnification x20, scale bar 20 µm. (B) Live imaging of Caco-2 cells fed with pHrodo-red zymosan over-night shows fluorescent internalize zymosan (red). Intact (arrowheads) and fragmented processed (arrow) zymosan are indicated. Original magnification x40, scale bar 20 µm. All images are representative of at least three independent experiments performed.

**Supp Fig. S3:**
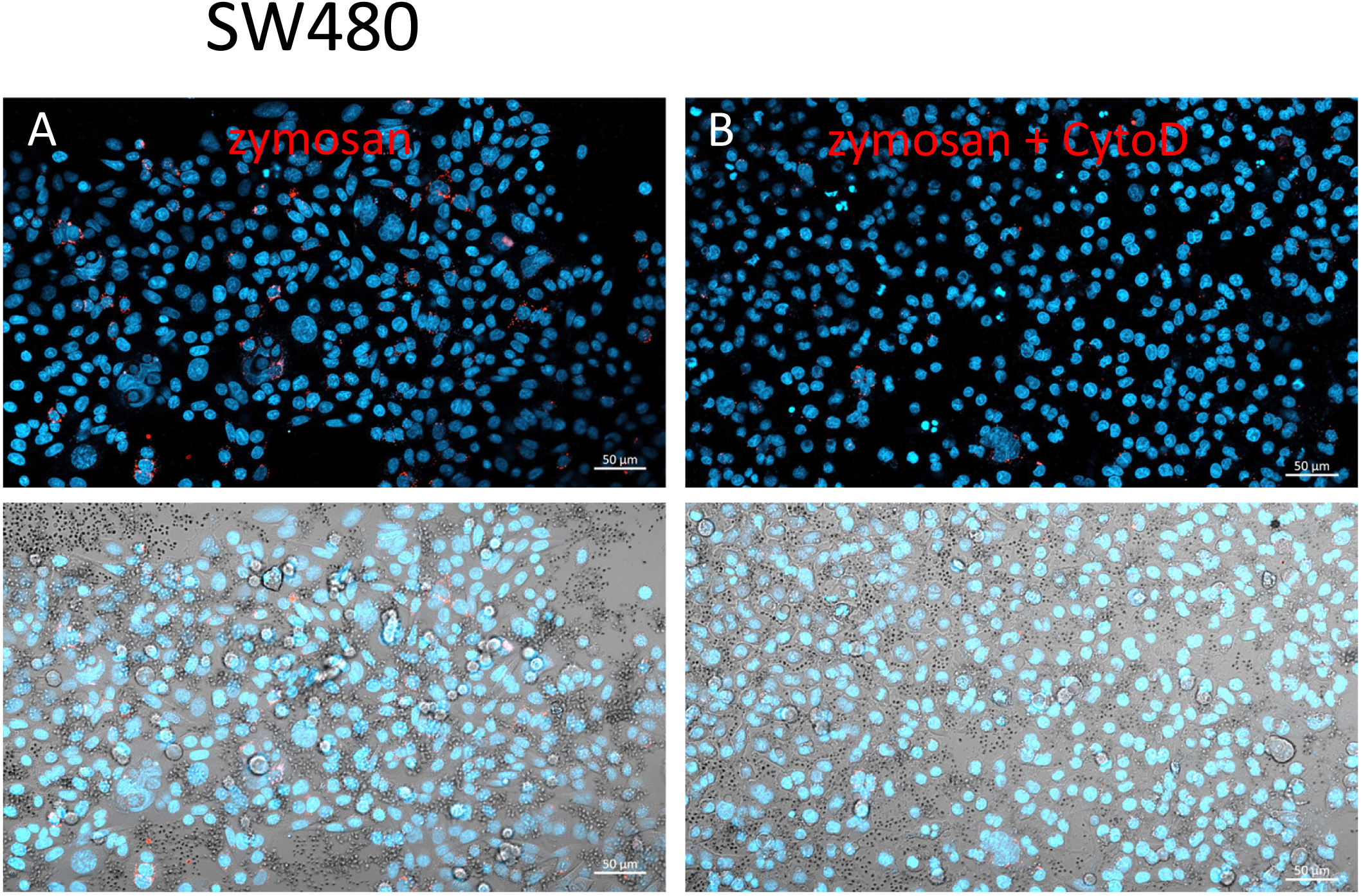
Wider fields of the images shown in Fig. 1 B-C. Zymosan uptake is sensitive to cytochalasin-D. SW480 cells were seeded on glass-bottom chambers as indicated in Methods, and fed over-night with pHrodo-red zymosan in the absence (A) or presence (B) of cytochalasin-D (10 µM) and counter stained with Hoechst 33342 (blue) prior to confocal live imaging. Original magnification x20, scale bar 10 µm.

**Supp Fig. S4:**
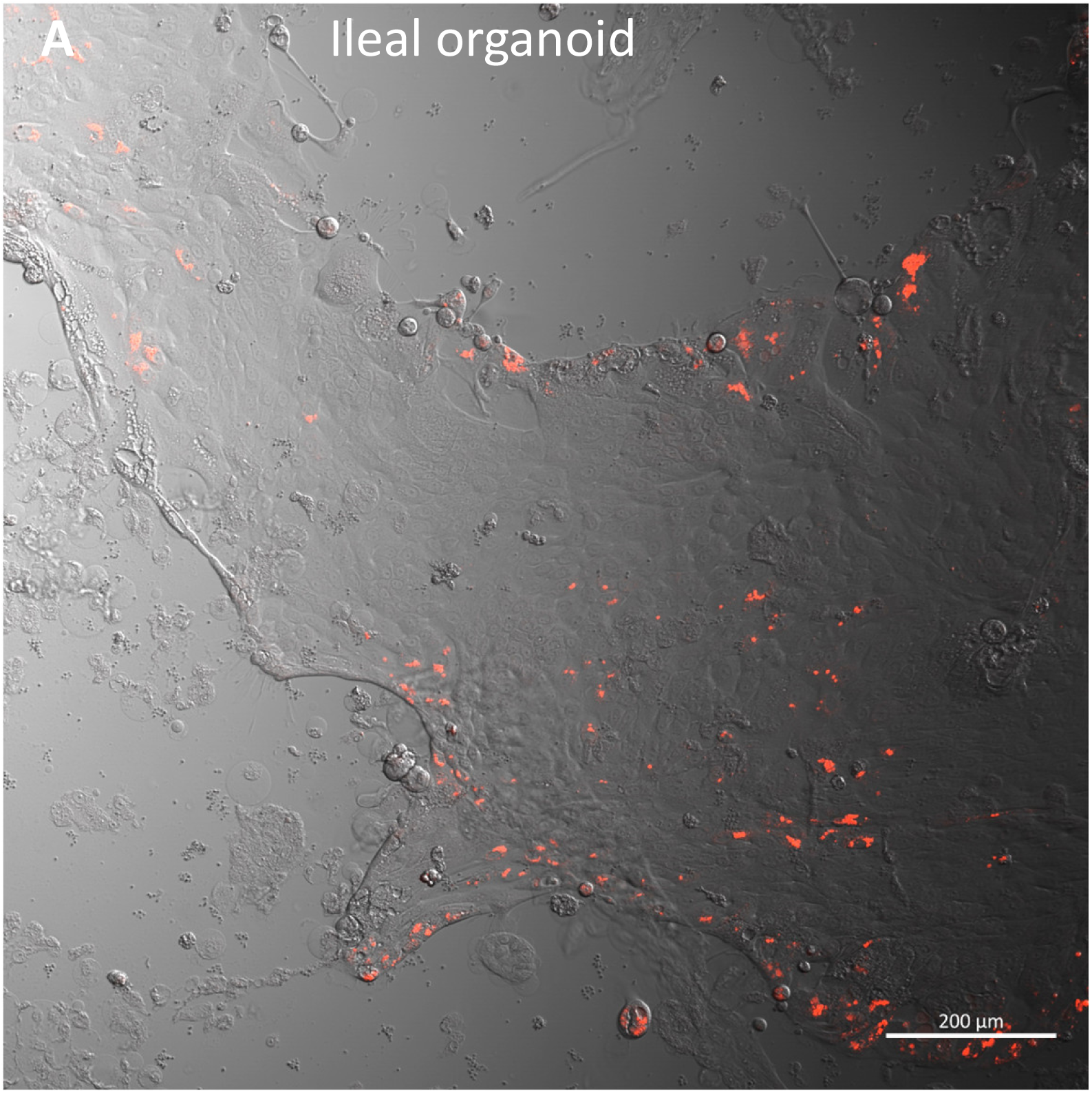
Phagocytosis of pHrodo-red zymosan by ileal organoids. Ileal organoids were grown as monolayers in expansion medium and let do differentiate for 2 days. PHrodo-red zymosan was added to the medium for 24h, and intracellular zymosan became red. Original magnification x10, scale bar 200 µm. Shown are representative images (from which a selected frame is shown in figure 2A), from 3-5 randomly acquired scans of 4 ileal and organoids.

**Supp Fig. S5:**
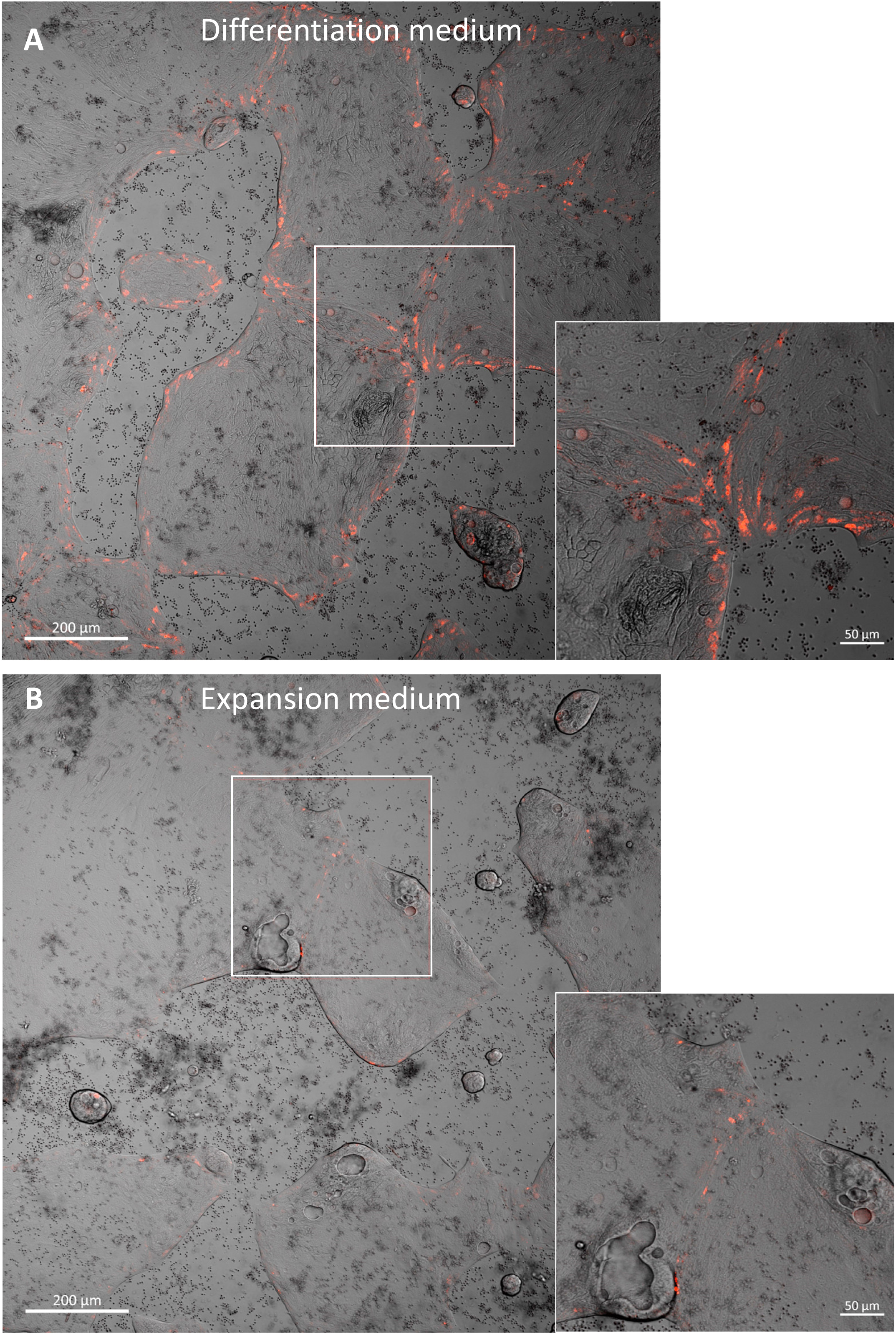
Phagocytosis of pHrodo-red zymosan by colonic organoids. Colonic organoids were grown as monolayers in expansion medium and let do differentiate for 2 days (A), or left in expansion medium (B). PHrodo-red zymosan (red) was added to the medium for 24h. Original magnification x10, scale bar 50 µm. Shown are representative images (from which selected frames (while squares) are enlarged in the insets), from 3-5 randomly acquired scans of two independent experiments of two organoids.

**Supp Fig. S6:**
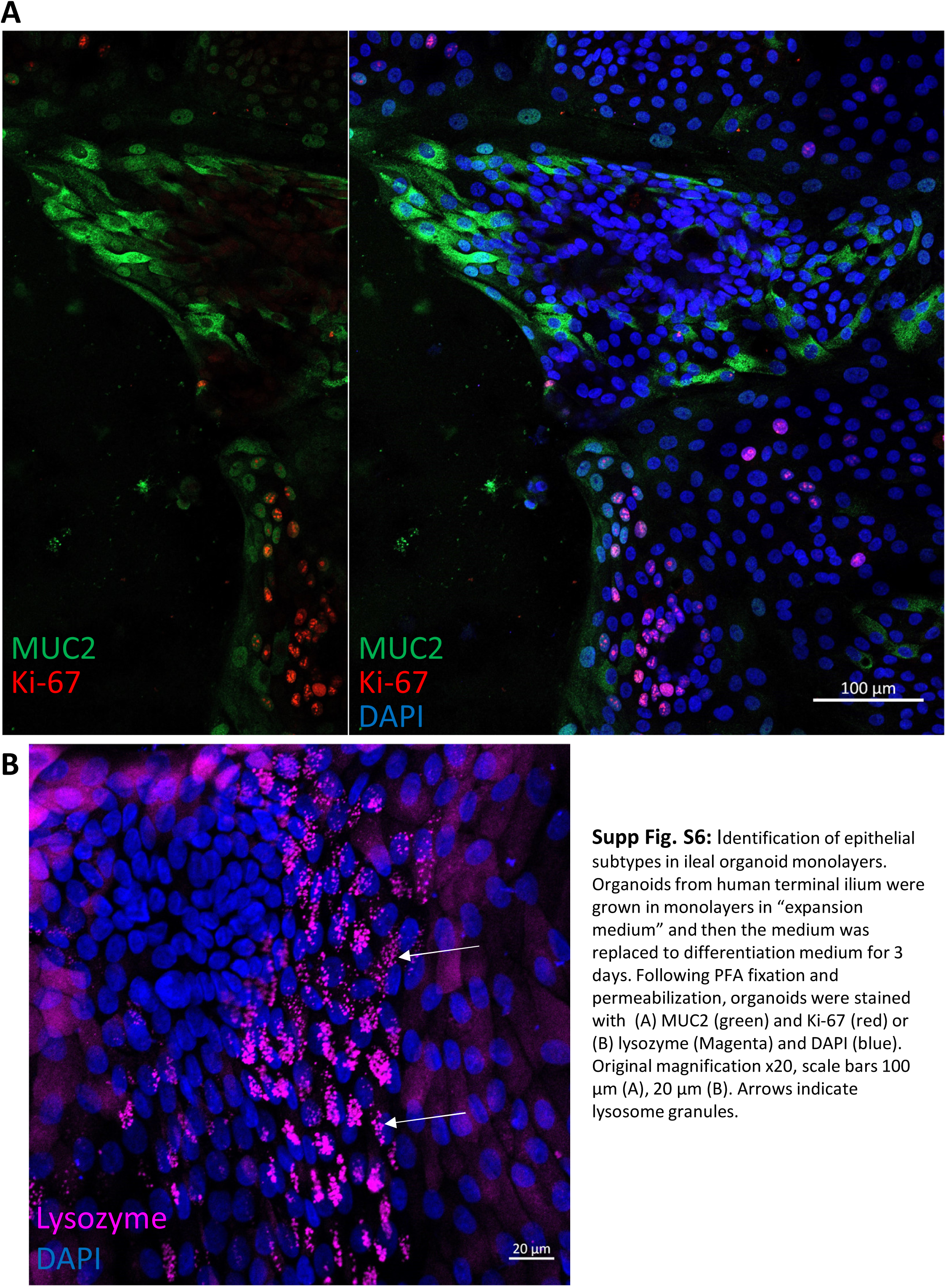
Identification of epithelial subtypes in ileal organoid monolayers. Organoids from human terminal ilium were grown in monolayers in “expansion medium” and then the medium was replaced to differentiation medium for 3 days. Following PFA fixation and permeabilization, organoids were stained with (A) Muc2 (green) and Ki-67 (red) or (B) lysozyme (Magenta) and DAPI (blue). Original magnification x20, scale bars 100 µm (A), 20 µm (B). Arrows indicate lysosome granules.

**Supp Fig. S7:**
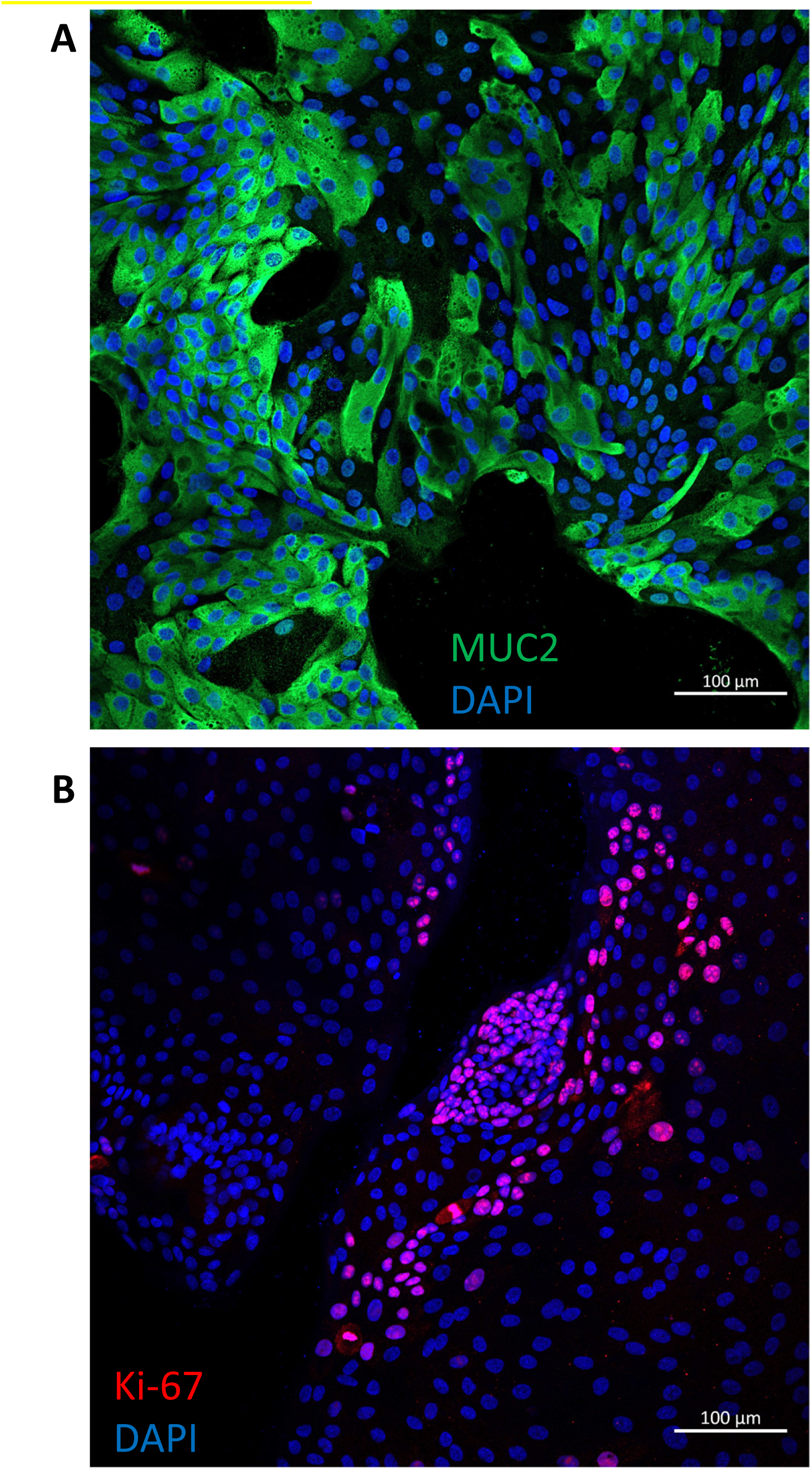
Identification of epithelial subtypes in colonic organoid monolayers. Organoids from human colon were grown in monolayers in “expansion medium” and then the medium was replaced to differentiation medium for 3 days. Following PFA fixation and permeabilization, organoids were stained with (A) MUC2 (green) and (B) Ki-67 (red) and DAPI (blue). Original magnification x20, scale bars 100 µm

**Supp Fig. S8:**
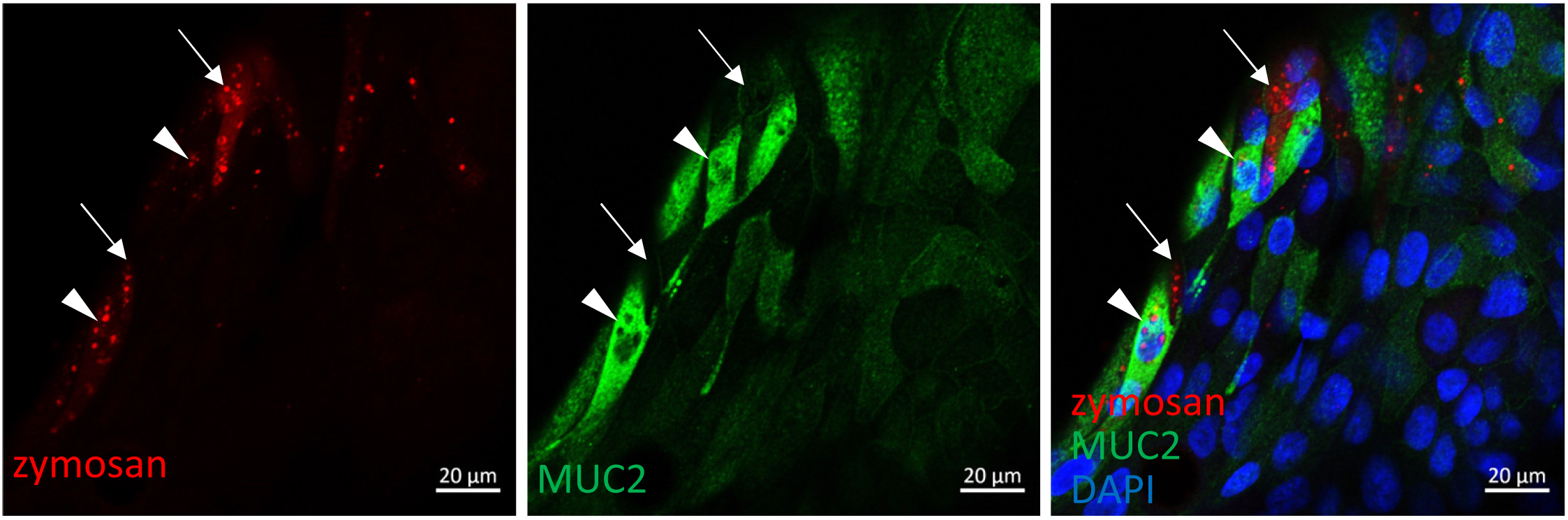
Zymosan is phagocytosed by goblet and non-goblet cells. Colonic organoids were fed over-night with pHrodo red zymosan (red), fixed and stained with MUC2 antibody (green) and DAPI (blue). Both goblet (MUC2+, arrowheads) and non-goblet (MUC2-, arrows) cells phagocytosed zymosan. Magnification x20, scale bar 20µm.

**Supp Fig. S9:**
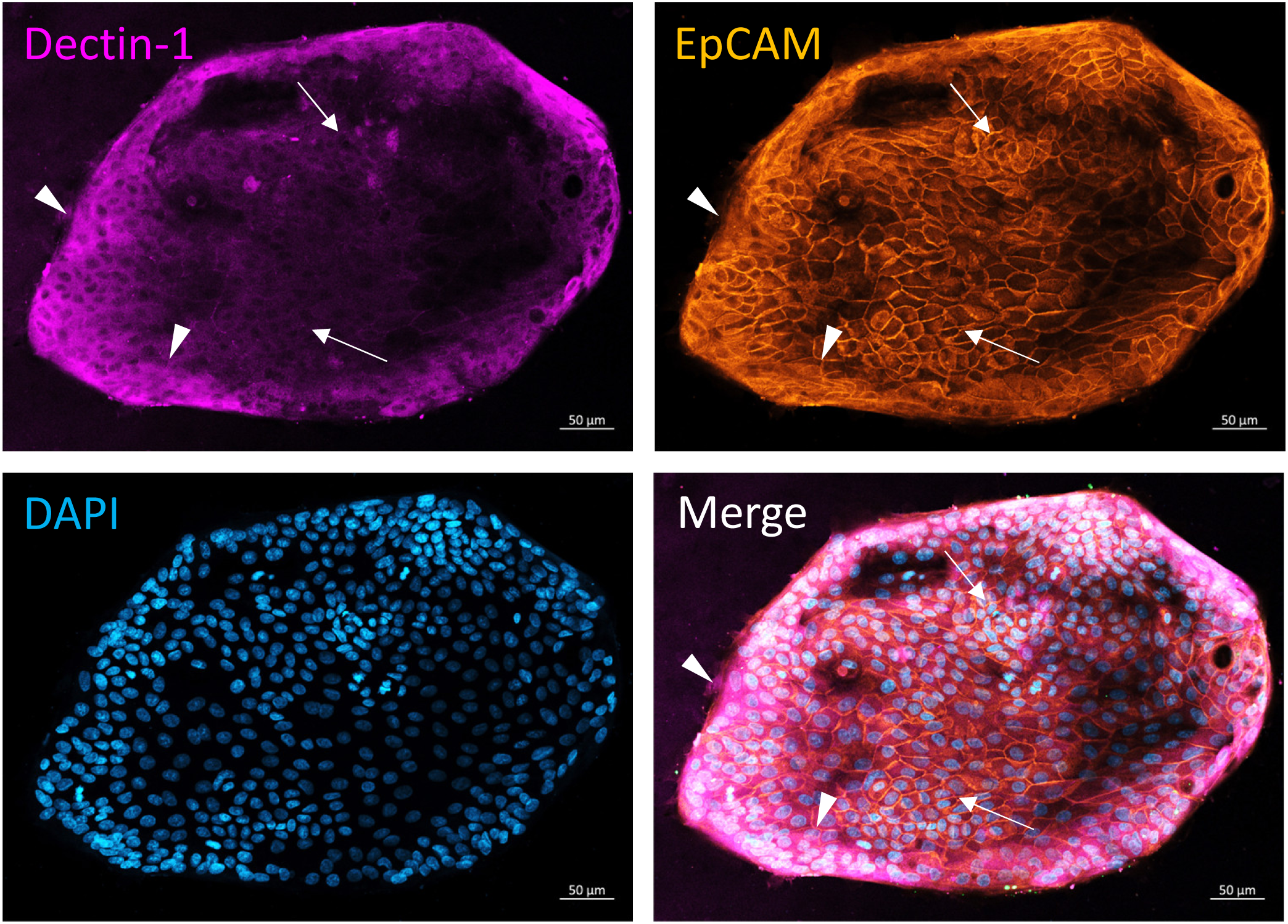
Dectin-1 is expressed mostly at the distal-peripheral region of colonic organoids. Colonic organoids were grown in monolayer and let to differentiate for 3 days. Following fixation with PFA, organoids were stained with Dectin-1 monoclonal antibody (magenta) and EpCAM polyclonal antibody (orange) and counterstained with DAPI (blue). While EpCAM staining is uniform in most of the organoid, Dectin-1 is enhanced mostly at the periphery of the organoid (arrow head), and less to the internal parts (arrows). Original magnification x20, scale bar 50 µm.

**Supp Fig. S10:**
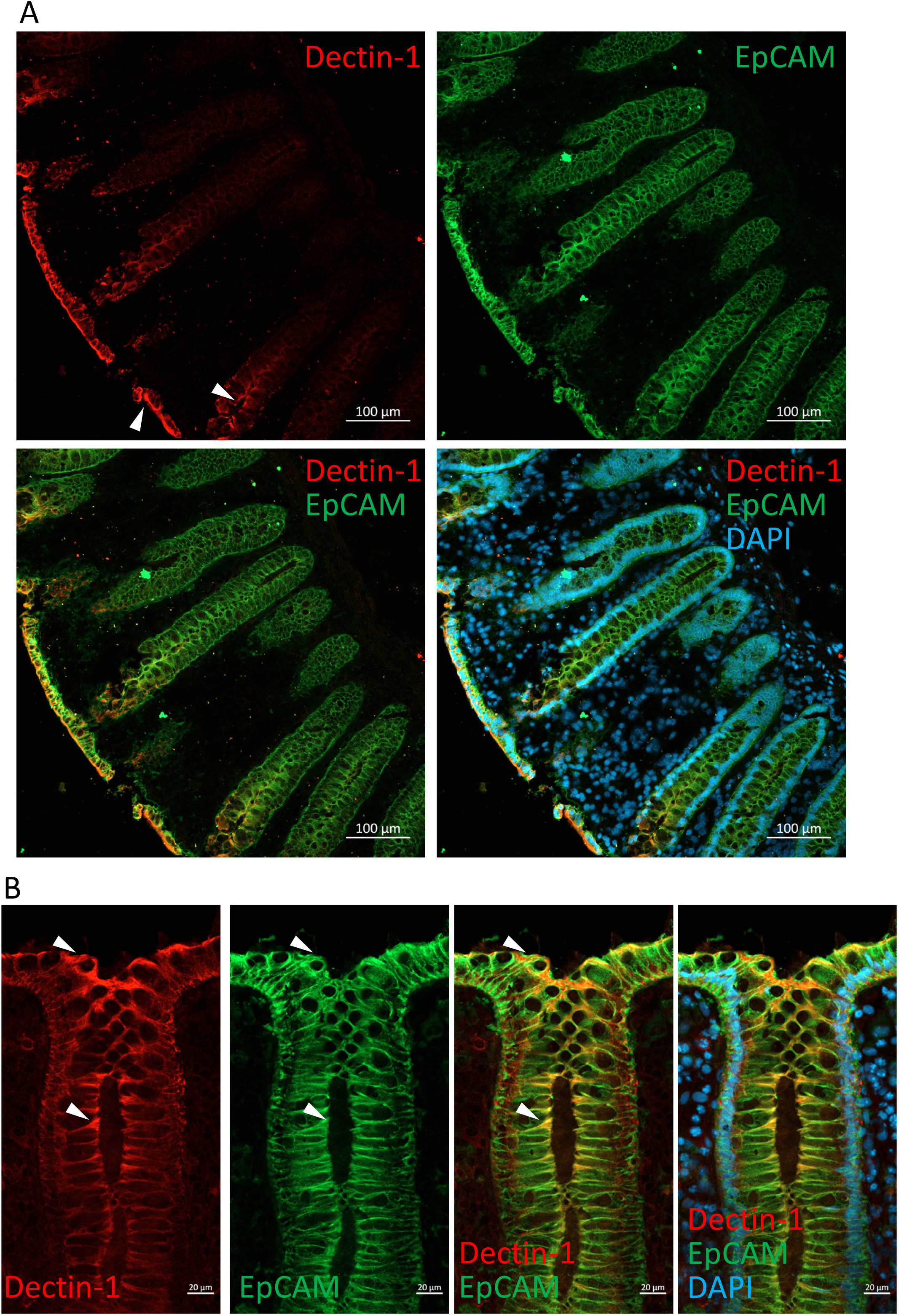
Dectin-1 is expressed in the luminal face of human intestinal crypts. (A, B) Frozen sections of human colon surgical specimen were stained with Dectin-1 polyclonal antibody (red) and EpCAM monoclonal antibody (green) and counterstained with DAPI (A, cyan). (A) shows that Dectin-1 is enhanced in the lumen facing IECs (arrowheads) while EpCAM stains uniformly IECs along the crypt, and (B) demonstrates that Dectin-1 is located to the apical face IECs (arrowheads). Original magnification x20, scale bar 100 µm(A) and 20 µm (B).

**Supp Fig. S11:**
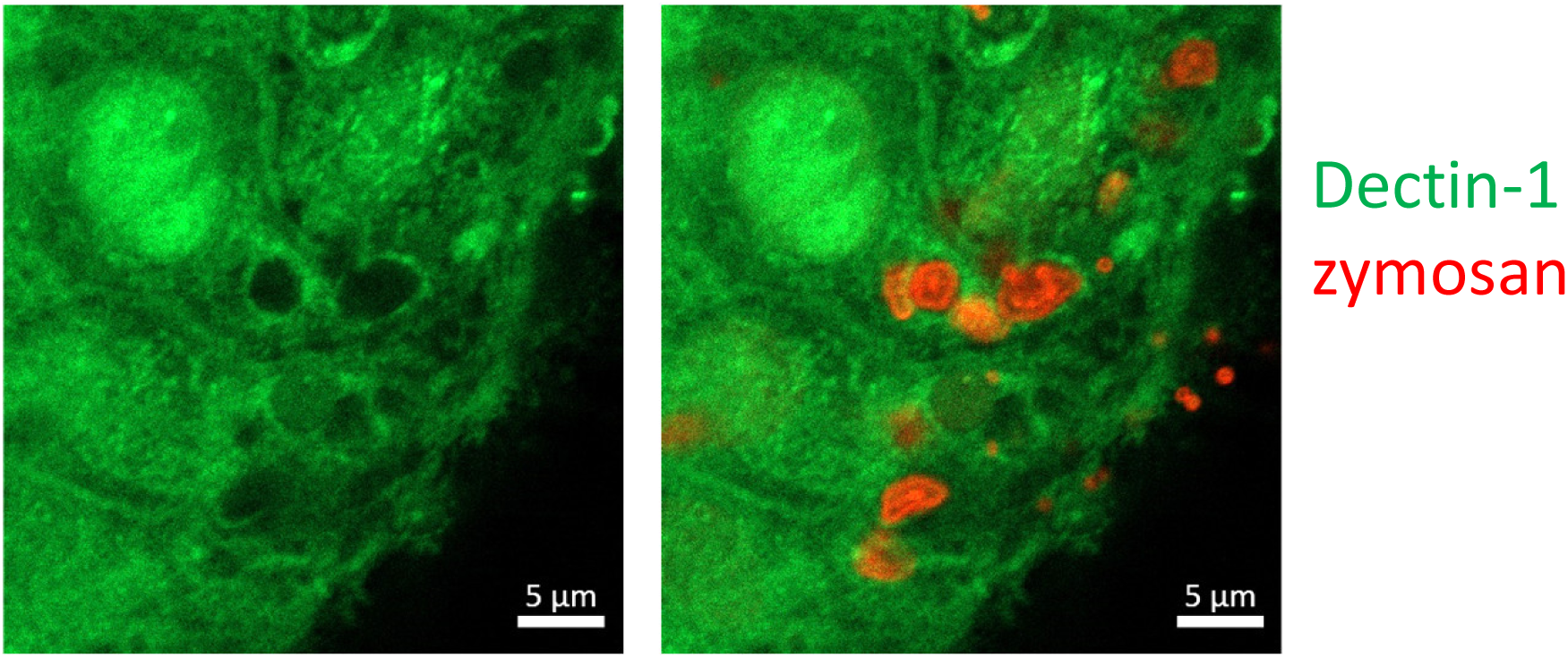
Dectin-1 is recruited to internalized zymosan. Colonic organoids were fed with pHrodo red zymosan and stained with FITC labeled Dectin-1 monoclonal antibody. Original magnification x63, scale bar 5µm.

**Supp Fig. S12:**
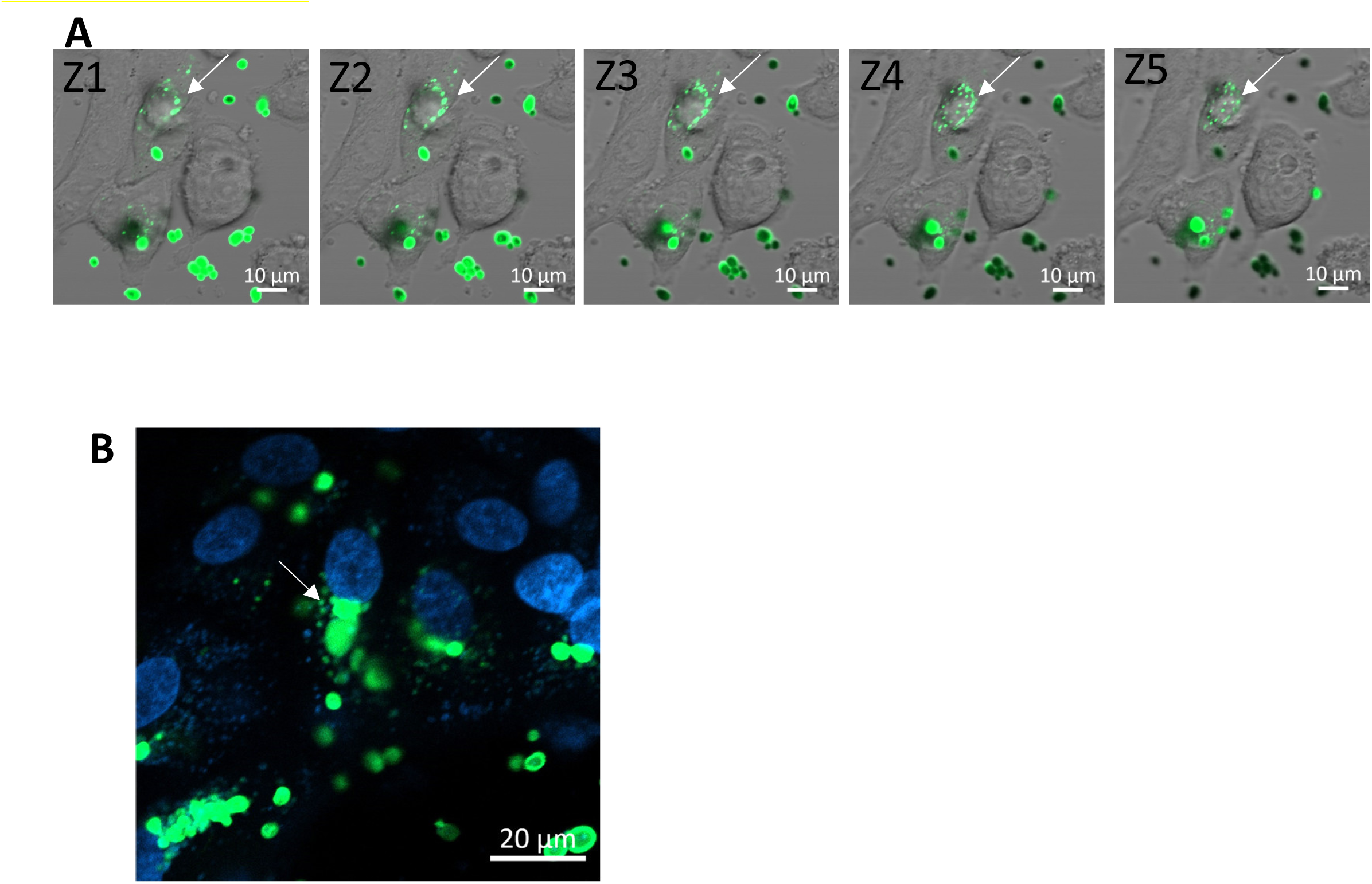
Phagocytosis of *C. albicans* by IECs. (A) SW480 cells were fed over-night with Rhodamine-green labeled HK *C. albicans*. Live images were taken (z-stuck 1 µm step). Arrow-intracellular fragmented *C. albicans*. (B) Ileal organoids were fed over-night with Rhodamine-green labeled UV-inactivated *C. albicans*, and stained with Hoechst 33342 prior to live confocal imaging. Arrow-intracellular fragmented *C. albicans*. Original magnification x20, scale bar 10 µm (A) 20 µm (B).

**Supp Fig. S13:**
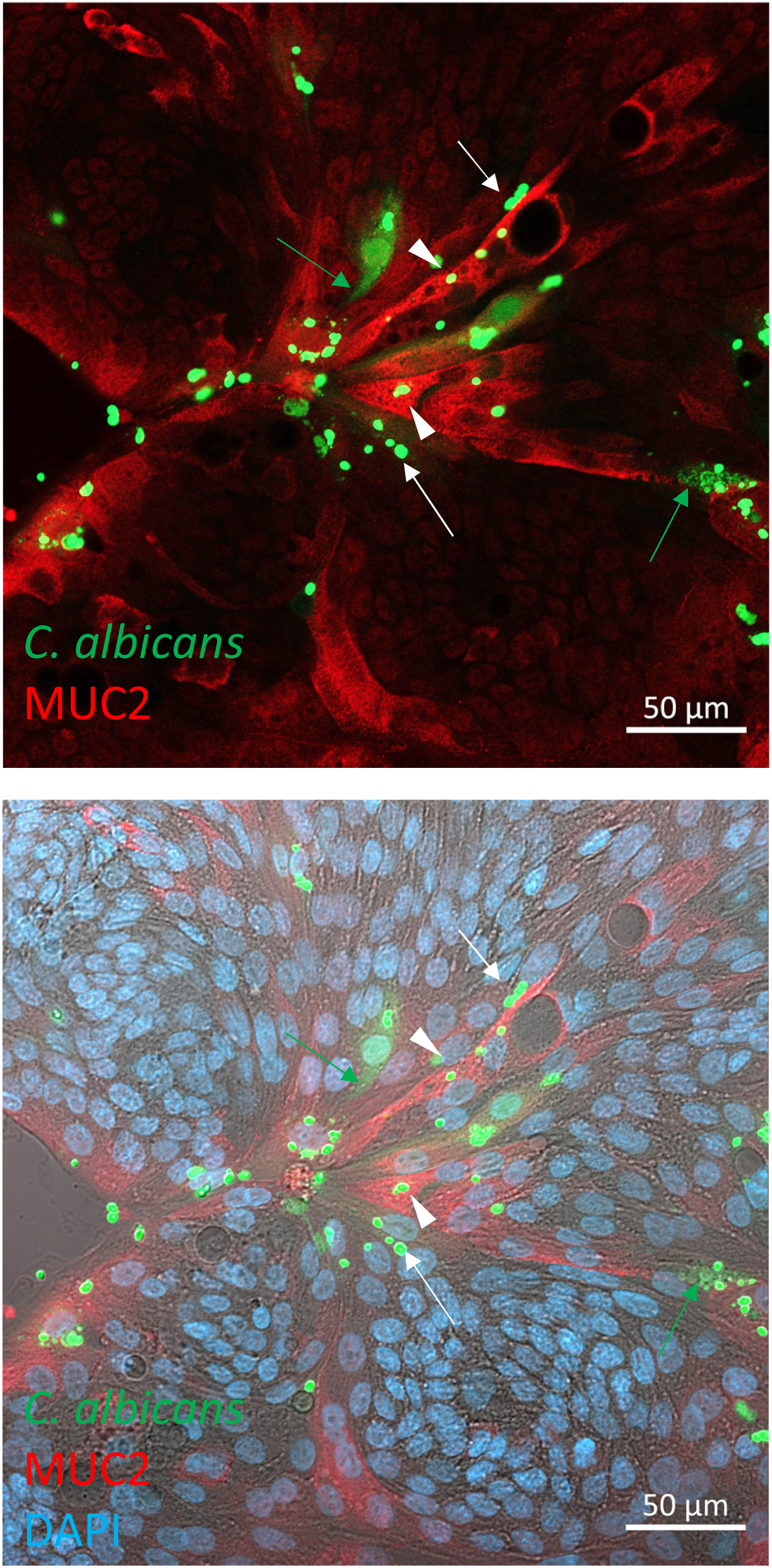
*C. albicans* is phagocytosed by goblet and non-goblet cells. Colonic organoids grown as monolayers were fed over-night with rhodamine-green labelled HK-*C. albicans* (green), fixed and stained with MUC2 antibody (red) and DAPI (blue). Both goblet (MUC2^+^, arrowheads) and non-goblet (MUC2^-^, arrows) cells phagocytosed *C. albicans*. Fragmented and diffused *C. albicans* (green arrows) indicate intracellular localization and processing. Magnification x20, scale bar 50 µm.

**Supp Fig. S14:**
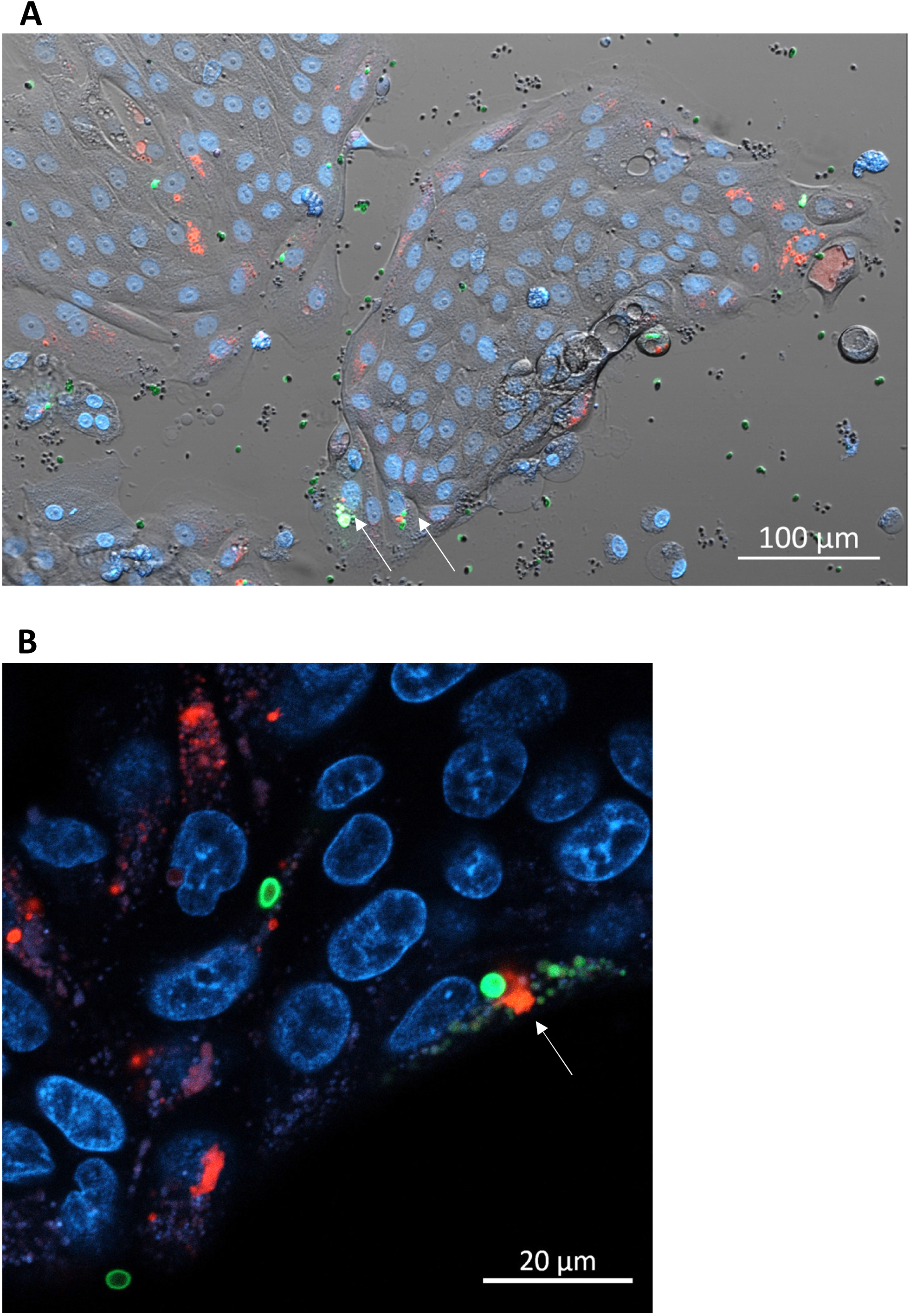
IECs can phagocytose both zymosan and *C. albicans*. Ileal (A) and colonic (B) organoids were fed over night with pHrodo-red zymosan and Rhodamine-green-X labeled HK-*C. albicans* and stained with Hoechst 33342 prior to confocal live imaging. Arrows indicate cells that phagocytosed both *zymosan and C. albicans.* Original magnification x10 (A) x40 (B), scale bar 100 µm (A) and 20 µm (B).

**Supp Fig. S15:**
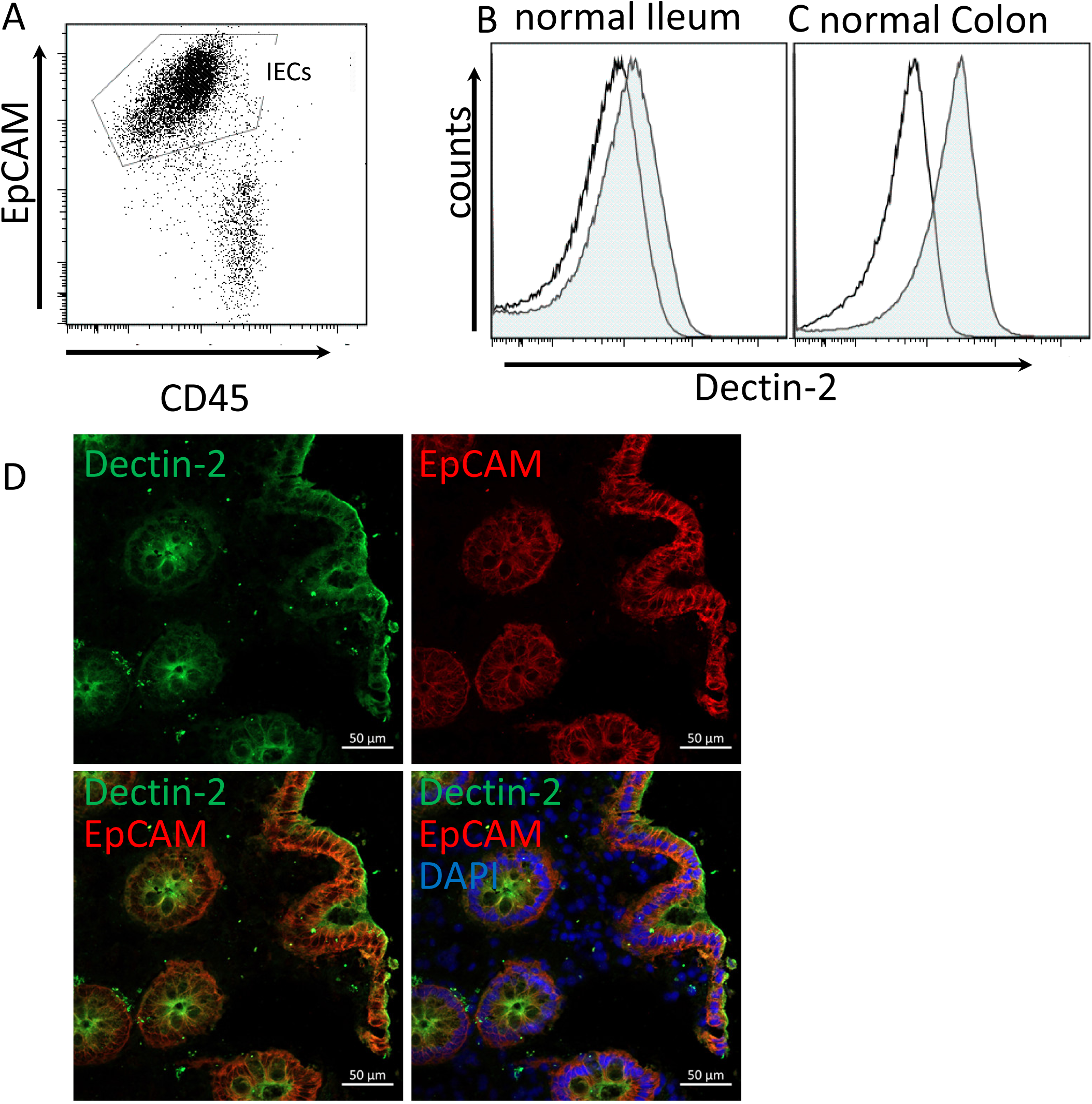
Dectin-2 is expressed on primary human IECs. (A) EpCAM-based flow cytometric gating on IECs. Freshly isolated human IEC-enriched preparation was stained with EpCAM and CD45 antibodies to identify IECs and leukocytes respectively. (B-C) Human IEC cell surface expression of Dectin-2. IECs generated from the (B) ileum and (C) colon of the same donor were stained as above, as well as with Dectin-2 Ab. Dectin-2 expression by intestinal epithelial cells (gated EpCAM+ population) was assessed by flow cytometry (filled gray histograms) and was compared with isotype-matched control Ab (open histograms). The data shown are representative of at least 3 individuals. (D) Frozen sections of human colon surgical specimen were stained with Dectin-2 monoclonal antibody (green) and EpCAM polyclonal antibody (red) and counterstained with DAPI (blue). Original magnification x20, scale bar 50 µm. The images are representative of three individuals tested.

**Supp Fig. S16:**
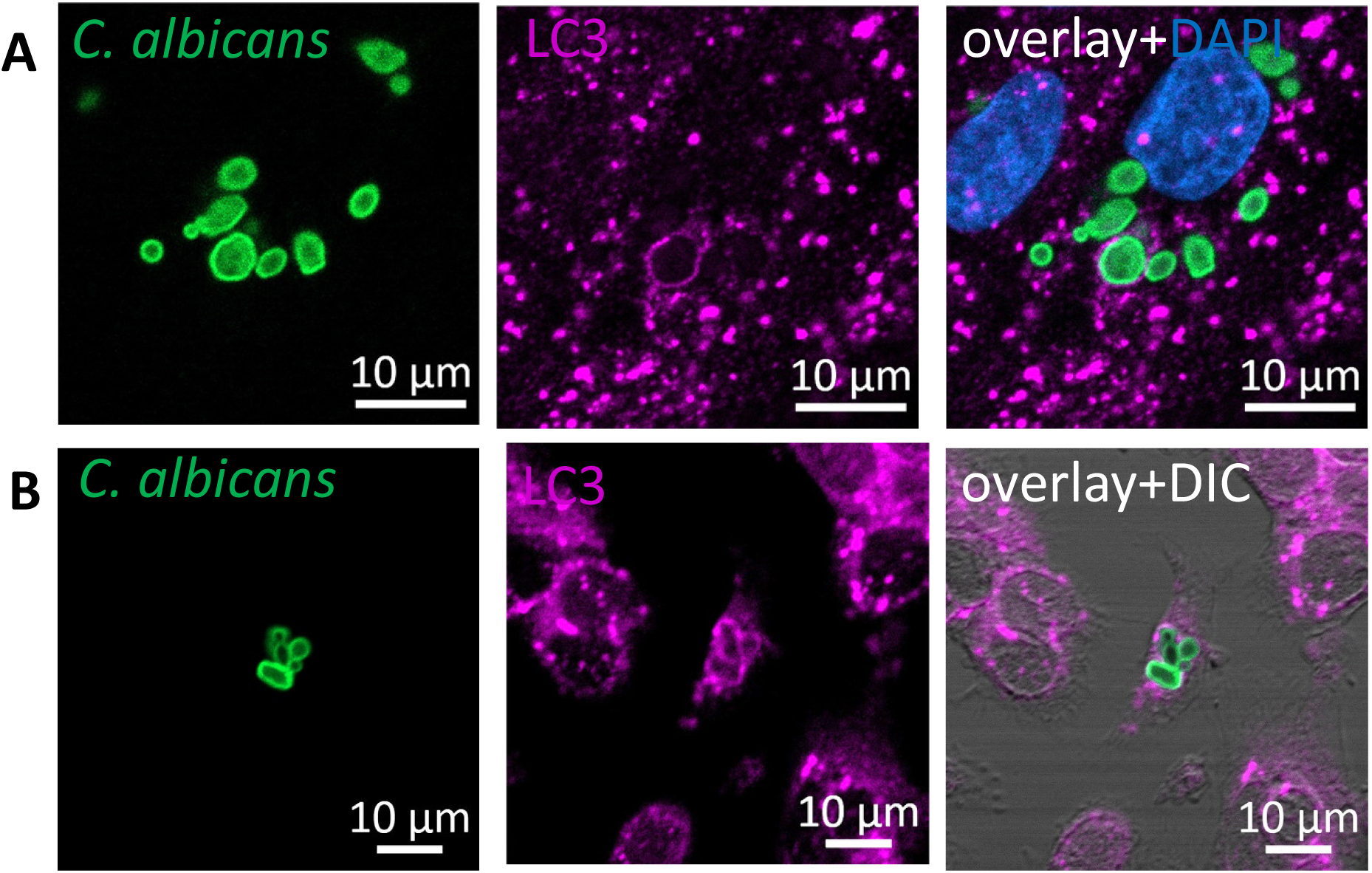
LC3 is recruited to phagocytosed particles. SW480 cells (A) and Ileal organoids (B) were fed with Rhodamine-green-X HK-*C. albicans* and stained with LC3 (magenta) and DAPI (blue). Original magnification x20 (A) x63 (B), scale bar 10 µm.

**Supp Fig. S17:**
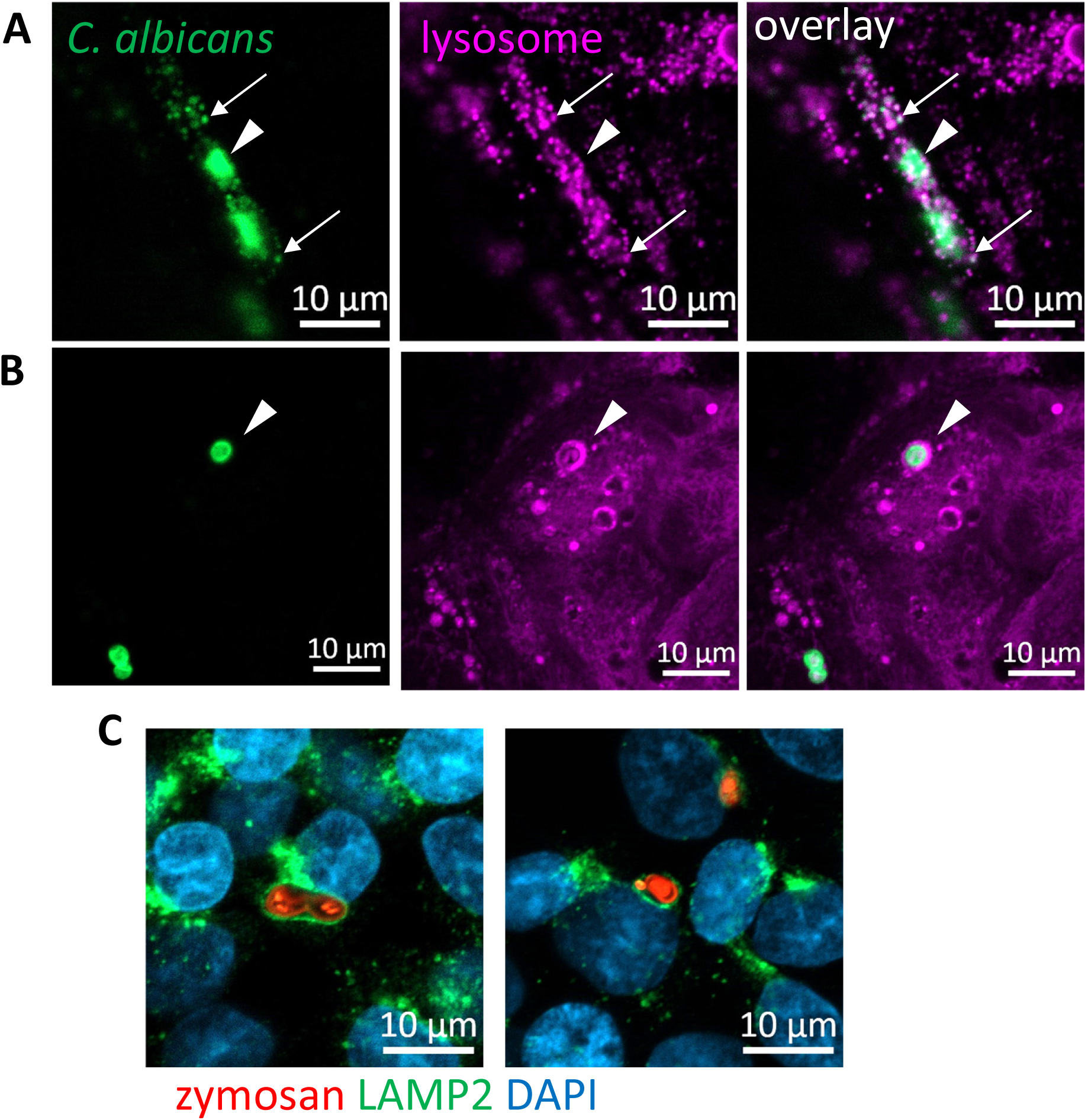
Phagocytosed particles are directed to the lysosomes. (A, B) Colonic organoids were fed with UV-inactivated *C. albicans* (green) and stained with lysosomal-NIR reagent (Magenta). Processed (A, arrow) and intact (A, B, arrowheads) particles are indicated, and co-localized with lysosomes. Original magnification x40 (A) x63 (B), scale bar 10 µm C) HCT116 cells were fed with AF594-zymosan (red) and stained with LAMP2 antibody (green) and DAPI (blue). Original magnification x63, scale bar 10 µm

**Supp Movie 1**: Zymosan uptake by intestinal organoids (related to Fig. 2C). differentiated monolayers of colonic organoids were fed with pHrodo-red zymosan for 24 hours. nuclei were stained with Hoechst 33342 (blue) prior to confocal live imaging. Shown a movie of the sequential frames acquired by z-stuck analysis. Original magnification x63, scale bar 5 µm.

**Supp. Movie 2**: Intestinal organoids uptake zymosan (related to Fig. 2C). Colonic organoids were treated as in Supp. Movie 1. Shown a movie of the sequential frames acquired by z-stuck analysis. Original magnification x63, scale bar 10 µm.

